# Minimal genetically encoded tags for fluorescent protein labeling in living neurons

**DOI:** 10.1101/2021.01.14.426692

**Authors:** Aleksandra Arsić, Cathleen Hagemann, Nevena Stajković, Timm Schubert, Ivana Nikić-Spiegel

## Abstract

Modern light microscopy, including super-resolution techniques, brought about a demand for small labeling tags that bring the fluorophore closer to the target. This challenge can be addressed by labeling unnatural amino acids (UAAs) with click chemistry. UAAs are site-specifically incorporated into a protein of interest by genetic code expansion. If the UAA carries a strained alkene or alkyne moiety it can be conjugated to a tetrazine-bearing fluorophore via a strain-promoted inverse-electron-demand Diels–Alder cycloaddition (SPIEDAC), a variant of bioorthogonal click chemistry. The minimal size of the incorporated tag and the possibility to couple the fluorophores directly to the protein of interest with single-residue precision make SPIEDAC live-cell labeling unique. However, until now, this type of labeling has not been used in complex, non-dividing cells, such as neurons. Using neurofilament light chain as a target protein, we established SPIEDAC labeling in living primary neurons and applied it for fixed-cell, live-cell, dual-color pulse—chase and super-resolution microscopy. We also show that SPIEDAC labeling can be combined with CRISPR/Cas9 genome engineering for tagging endogenous NFL. Due to its versatile nature and compatibility with advanced microscopy techniques, we anticipate that SPIEDAC labeling will contribute to novel discoveries in neurobiology.

## Introduction

Fluorescence microscopy encompasses experimental techniques that are central to the study of cellular and molecular (neuro)biology. These techniques allow the investigation of protein localization and interactions, as well as the visualization of protein dynamics in real time. The ability to investigate these processes in living cells and with high resolution is especially relevant for polarized cells, such as neurons. Aggregation of proteins and perturbation of protein processing, turnover and axonal transport mechanisms, have been implicated in a number of neurological diseases. Advanced light microscopy techniques, including live-cell and super-resolution imaging are indispensable for elucidating the underlying molecular mechanisms, but to fully utilize their potential, advances in imaging technologies need to be matched with advancement in the field of protein labeling^1–5^.

Breaking the resolution limit in modern super-resolution microscopy techniques has brought about a demand for small labeling tags that bring the fluorophore close to the target. Consequently, there is a trend towards using smaller nanobodies, affimers, and aptamers instead of the conventional primary–secondary antibody complexes^6–8^. Similarly, in live-cell labeling, relatively large fluorescent proteins (FPs) are being substituted with small fluorescent dyes^9,10^. The reasons are plentiful: fluorescent organic dyes are not only 20-fold smaller (approx. 0.5–2k Da compared to 25 kDa for GFP) but they can also offer superior photophysical properties and more spectral variants compared to FPs. Consequently, a number of methods for the attachment of fluorescent dyes to proteins of interest have been developed, such as widely used Halo^11^, SNAP^12^, and CLIP tags^13^. However, similarly to genetically encoded FPs, these labeling approaches require making protein fusions, which can affect the function of the protein of interest (POI). This is a general problem that also applies to neuroscience studies involving neuronal cytoskeletal elements, ion channels, receptors and other synaptic proteins^14^.

As an alternative, site-specific labeling utilizing unnatural amino acids (UAAs, also referred to as noncanonical or non-natural amino acids) and bioorthogonal click chemistry is emerging as a powerful approach for directly labeling proteins with small organic fluorophores^15^. These reactions are perhaps best exemplified by the strain-promoted inverse electron-demand Diels–Alder cycloaddition (SPIEDAC) between a strained alkene/alkyne and a tetrazine^16,17^. Due to its extremely high reactivity (reaction rates >10^5^ M^-1^ s^-1^) and live-cell compatibility, SPIEDAC is a perfect candidate for the fast, specific and efficient labeling of biomolecules. To exploit these advantages for protein labeling in living systems, UAAs carrying click-chemistry-reactive moieties in their side chains (“clickable” UAAs) need to be incorporated into target proteins. This can be achieved by amber codon suppression, an emerging area of genetic code expansion^18,19^ utilizing orthogonal translational machinery to direct co-translational and site-specific UAA incorporation into the POI (**Fig. 1a**). SPIEDAC-reactive UAAs, such as TCO‪A-Lys (**Fig. 1b**), contain strained alkyne or alkene moieties in their side chains. They can be genetically encoded in mammalian cells with the help of the naturally occurring pyrrolysine (Pyl)-specific amber codon suppressor tRNA^Pyl^ and its cognate aminoacyl-tRNA synthetase (PylRS) from *Methanosarcina* species (*M. barkeri* and *M. mazei)*, the binding pocket of which has been modified to accommodate the side chains of clickable UAAs^20–23^. SPIEDAC reactions between genetically encoded clickable UAAs and tetrazine derivatives of fluorescent dyes combines a fast, bioorthogonal reaction with site-specific labeling (**Fig. 1a, b**). This type of labeling provides several advantages, the most notable being the opportunity to attach the fluorescent dye with single-residue precision directly to the POI. This reduces the potential negative impact of the fluorescent tag on the function of the POI and has steric advantages in the context of super-resolution imaging, as is discussed below.

**Fig 1.**
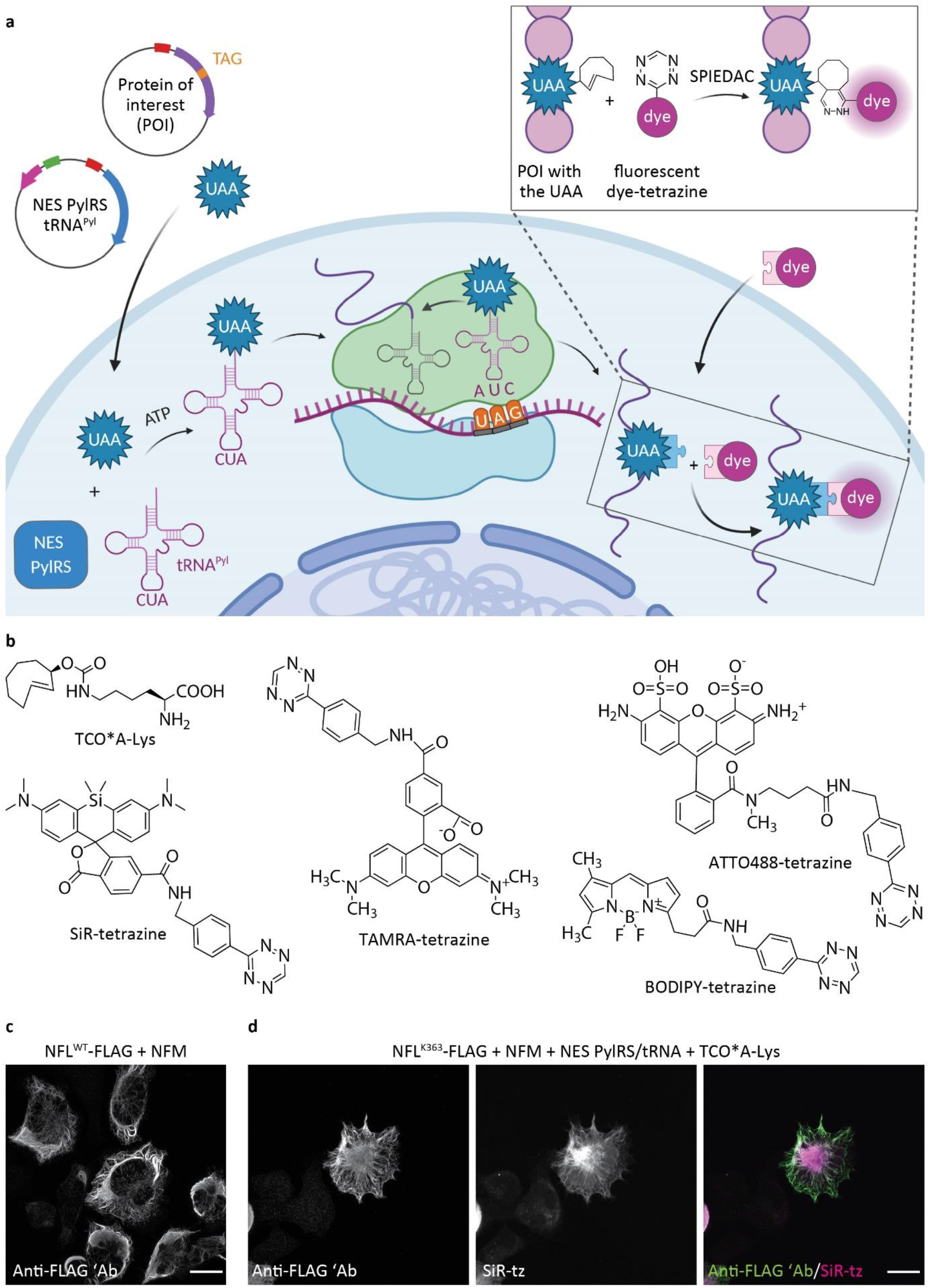
Genetic code expansion and click labeling of neurofilament light chain (NFL) in ND7/23 neuroblastoma cells. **a**, A schematic representation of genetic code expansion and labeling of proteins by click chemistry. Cells are transfected with plasmids bearing genes that encode for NES PylRS (a pyrrolysyl-tRNA synthetase with a nuclear export signal) and tRNA_CUA_^Pyl^, as well as a TAG stop codon-containing gene that encodes for the protein of interest (POI). The NES PylRS charges tRNA_CUA_^Pyl^ with the unnatural amino acid (UAA), this charged tRNA_CUA_^Pyl^ recognizes the UAG stop codon in the mRNA encoding the POI, and UAA is co-translationally incorporated into the POI. In the subsequent step, a tetrazine derivative of a fluorescent dye reacts with the UAA by strain-promoted inverse electron-demand Diels-Alder cycloaddition (SPIEDAC) reaction, and the dye is attached directly to the POI. **b**, Chemical structures of UAAs and tetrazine dyes that were used to obtain the data shown in the main figures. **c**, ND7/23 cells expressing NFM, NES PylRS/tRNA_CUA_^Pyl^, and wild-type NFL^WT^-FLAG, stained with an anti-FLAG antibody, followed by Alexa Fluor (AF) 488-conjugated secondary antibody. **d**, ND7/23 cells expressing NFL^K363TAG^-FLAG, neurofilament medium chain (NFM), and NES PylRS/tRNA_CUA_^Pyl^. Cells were incubated overnight with TCO‪A-Lys and then click labeled with silicon rhodamine-tetrazine (SiR-tz). Afterwards, cells were fixed and stained with the anti-FLAG antibody, followed by AF488-conjugated secondary antibody. *Z*-stack images were acquired with a confocal scanning microscope, and are shown as maximum intensity projections. Scale bars: 20 µm (**c**,**d**).

Since pioneering studies were published demonstrating its suitability for conventional imaging of epidermal growth factor receptor^20^ and super-resolution imaging of insulin receptor and influenza virus hemagglutinin protein^24^, SPIEDAC-based labeling has emerged as a powerful approach for protein labeling in living mammalian cells. In the meantime, amber codon suppression using a conventional PylRS/tRNA^Pyl^ pair, a more efficient PylRS variant fused to a nuclear export signal (NES PylRS)^25^, as well as rationally designed tRNA^Pyl^ variants^26^, has been applied for the fluorescent labeling and imaging of receptors, cytoskeletal proteins, nucleoporins, viral proteins, and other protein structures, as recently reviewed^27^. However, all of these studies were performed in readily transfected cell lines, and this type of labeling has never been used for microscopy studies in complex cells, such as primary neurons. Here, by using neurofilament light chain (NFL) as a target protein, we established UAA-based SPIEDAC-labeling in living neurons. The versatility of this labeling approach allowed us to use it for fixed-cell, live-cell, dual-color pulse–chase and super-resolution imaging of NFL. We also show that amber codon suppression and SPIEDAC labeling can be combined with CRISPR/Cas9 genome engineering for labeling of endogenous neuronal proteins.

## Results

### Establishing and optimizing genetic code expansion and click labeling in a rodent ND7/23 neuroblastoma cell line

To establish site-specific SPIEDAC-based fluorescent labeling of NFL (hereafter also referred to as “click labeling”) in primary neurons, we needed to identify optimal transfection conditions for wild-type NFL (NFL^WT^) and its amber mutants (NFL^TAG^). Proper expression of NFL^WT^ is a prerequisite for optimization of the subsequent click labeling step, which involved testing several NFL^TAG^ mutants and selecting the one that performed best in terms of expression and labeling efficiency. Because the transfection efficiency in primary neurons is very low, we first wanted to identify an intermediate host cell line for the amber codon suppression of NFL^TAG^ mutants. For this purpose, we tested ND7/23 cells, a neuron-derived immortalized cell line. NFL^WT^ expresses well in ND7/23 cells (**Fig. 1c**) and forms a neurofilament network when co-transfected with neurofilament medium chain (NFM).

To establish the click labeling of NFL^TAG^ in ND7/23 cells, we initially generated four different amber mutants: NFL^K211TAG^, NFL^K363TAG^, NFL^R438TAG^, and NFL^K468TAG^. When selecting the positions of the amber mutations, we avoided all residues known to undergo posttranslational modifications, as well as all those involved in the pathology of NFL-associated diseases. The creation of multiple site-specific TAG mutants is an essential step in establishing a protocol for click labeling a new target protein^28^. The reason for this is inherent to the amber codon suppression technology, the efficiency of which depends on the position and its nucleotide sequence context. In addition, the chosen TAG position needs to be specifically suppressed only in the presence of the UAA and should be accessible to a tetrazine dye for efficient SPIEDAC labeling. We initially tested silicon rhodamine (SiR)-tetrazine, a cell-permeable fluorophore that is widely used in live-cell and super-resolution microscopy studies^29^. The NFL^TAG^ mutants varied in their expression level and labeling efficiency when cells were co-transfected with the NESPylRS/tRNA^Pyl^ plasmid in the presence of the TCO‪A-Lys (**Supplementary Fig. 1**). Furthermore, screening of multiple mutants allowed us to identify and exclude those that show UAA-independent read-through of the TAG codon. For example, NFL^K211TAG^ exhibited an anti-FLAG signal in the absence of the UAA, suggesting synthesis of the full-length protein (**Supplementary Fig. 1a**); however, as was to be expected, labeling with SiR-tetrazine was unsuccessful. The mutant NFL^R438TAG^ was unable to form a normal neurofilament network and aggregated in the cytoplasm, indicating that arginine at this position might be involved in neurofilament interactions. Of the two other mutants, NFL^K363TAG^ was expressed at higher levels and was more efficiently labeled with SiR-tetrazine (**Fig. 1d** and **Supplementary Fig. 1b,d**). We subsequently used this mutant to compare different UAAs and other tetrazine dyes. In addition to TCO‪A-Lys, endo-BCN-Lys and TCO4en-Lys were genetically encoded into NFL^K363TAG^ and specifically labeled with SiR-tetrazine (**Supplementary Fig. 2**). We also evaluated other dyes for click labeling of NFL^K363TAG^ in living and fixed ND7/23 cells (**Supplementary Fig. 3**). Although click labeling of NFL^K363TAG^ was possible with both cell-permeable (SiR-tetrazine, Janelia Fluor (JF) 646-methyl-tetrazine, CF650-methyl-tetrazine, TAMRA-tetrazine, JF549-tetrazine, CF500-methyl-tetrazine, BODIPY-tetrazine) and cell-impermeable tetrazine dyes (Alexa Fluor 647-tetrazine, ATTO655-methyl-tetrazine, ATTO488-tetrazine), we noted that the cell-impermeable dyes gave higher background fluorescence, which was most likely caused by the fact that the labeling was done post-fixation and not in living cells. Taken together, these results show that ND7/23 neuroblastoma cells are a good host for the genetic code expansion and click labeling. They are transfected with high efficiency and allowed us to identify the most suitable NFL^TAG^ mutant and to optimize SPIEDAC labeling conditions.

**Fig 2.**
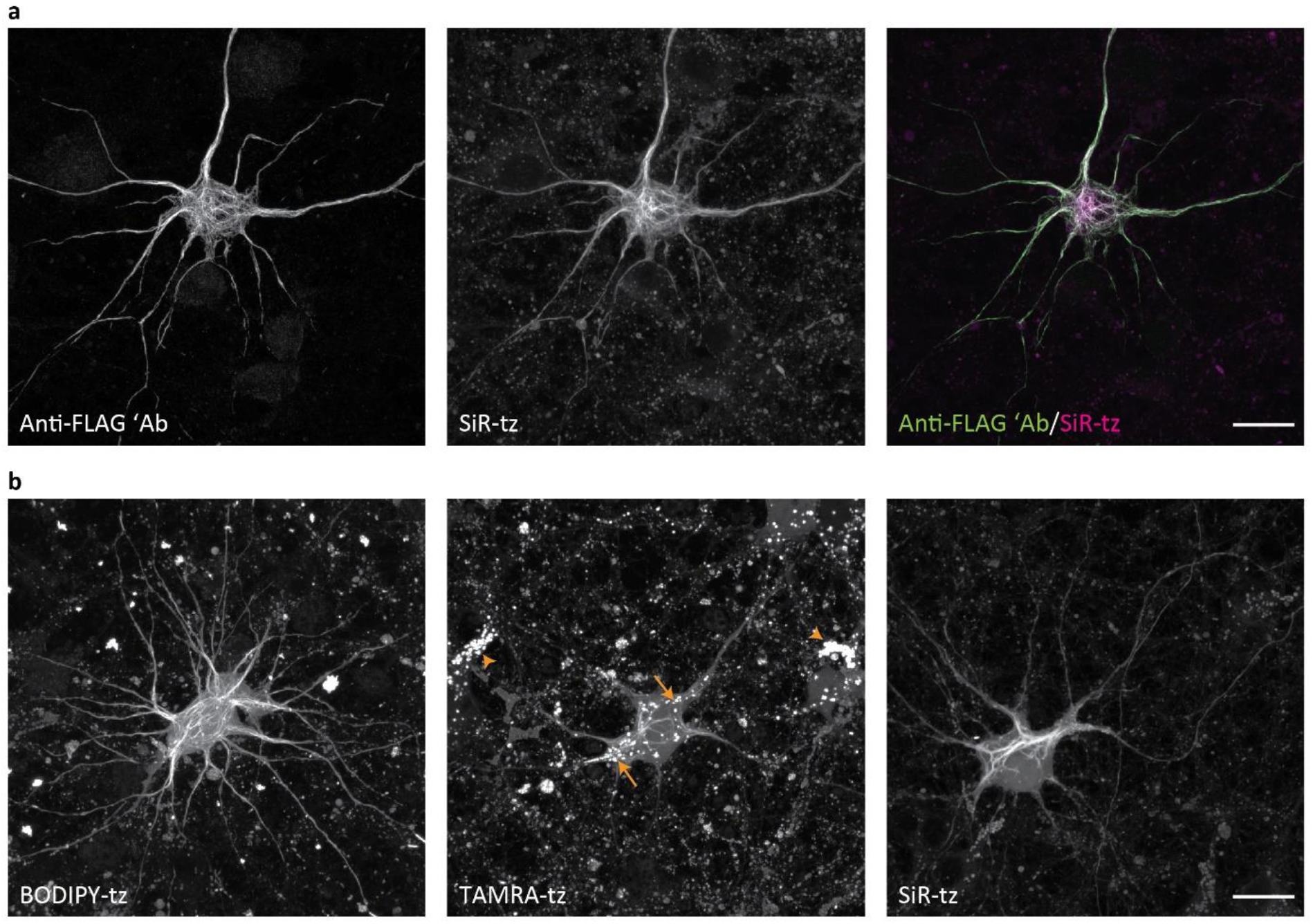
Genetic code expansion and click labeling of NFL in live primary mouse cortical neurons (MCNs). **a**,**b**, MCNs expressing NFL^K363TAG^-FLAG, NFM and NES PylRS/tRNA_CUA_^Pyl^ in the presence of TCO‪A-Lys. **a**, Three days after transfection, neurons were labeled with SiR-tz via click chemistry, then fixed and stained with anti-FLAG antibody, followed by AF488-conjugated secondary antibody. After fixation and immunocytochemistry labeling, neurons were imaged on a confocal scanning microscope. **b**, Three days after transfection, neurons were labeled with either BODIPY-tz, TAMRA-tz, or SiR-tz. Excess dye was washed for 3 h, and live neurons were imaged on a confocal scanning microscope. In TAMRA-tz panel, background staining of lysosomes is highlighted in both NFL^K363TAG^-expressing neuron (arrows) and in non-transfected neurons (arrowheads). *Z*-stack images are shown as maximum intensity projections. Scale bars: 20 µm (**a**,**b**).

**Fig 3.**
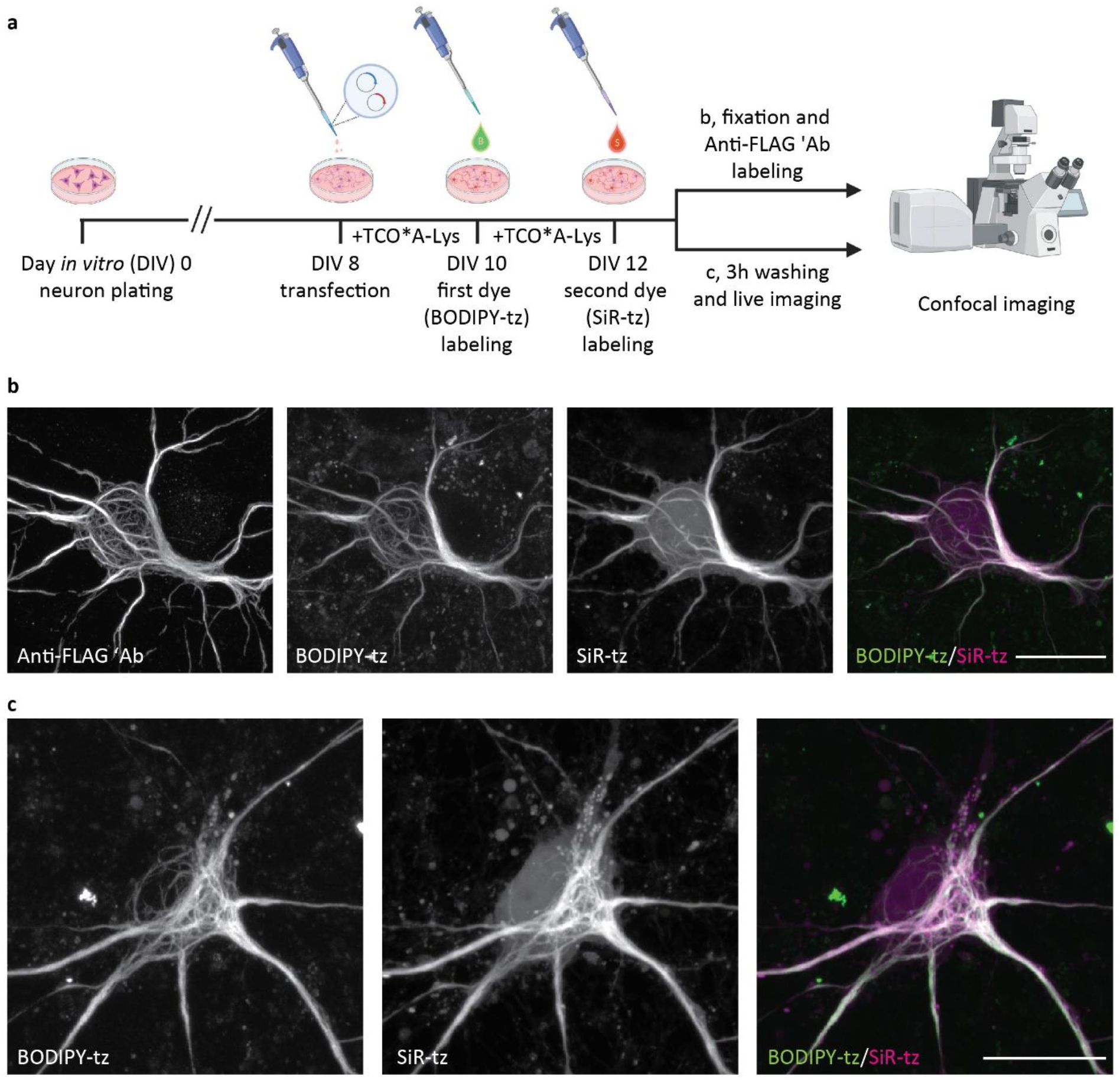
Pulse–chase click labeling of two NFL populations in live primary mouse cortical neurons (MCNs). **a**, A schematic representation of the experimental workflow. Eight days after plating, MCNs were transfected with NFL^K363TAG^-FLAG, NFM, and NES PylRS/tRNA_CUA_^Pyl^ constructs. After 2 days of incubation with TCO‪A-Lys, neurons were labeled with the first dye (BODIPY-tz), incubated with TCO‪A-Lys for a further 2 days, and labeled with the second dye (SiR-tz). After the second labeling step, neurons were either fixed, stained with anti-FLAG antibody followed by AF555-conjugated secondary antibody, and imaged on a confocal scanning microscope (**b**), or live neurons were imaged on a confocal scanning microscope (**c**). *Z-*stack images are shown as maximum intensity projections. Scale bars: 20 µm (**b**,**c**).

### Expression and click labeling of NFL^K363TAG^ in living primary mouse neurons

After identifying the optimal amber mutant position and conditions for click labeling of NFL^TAG^ in ND7/23 cells, we focused on experiments in primary mouse cortical neurons. When co-transfected with NESPylRS/tRNA^Pyl^ in the presence of TCO‪A-Lys, the NFL^K363TAG^ mutant was expressed well and formed neurofilament networks in primary neurons (**Fig. 2a**). Furthermore, an anti-FLAG antibody co-localized with the SiR-tetrazine signal, indicating that specific click labeling of NFL^K363TAG→TCO‪A-Lys^ had occurred (**Fig. 2a**). In addition to SiR-tetrazine, we evaluated the suitability of other cell-permeable and cell-impermeable dyes for click labeling of NFL^K363TAG→TCO‪A-Lys^ in living and fixed neurons, respectively (**Supplementary Fig. 4**). NFL^K363TAG→UAA^ showed higher expression levels in neurons than in ND7/23 cells, and consequently higher intensity of click-labeled NFL. This is exemplified by JF646-methyl-tetrazine, which gave a weak signal in ND7/23 cells after the labeling step compared to that observed in primary neurons (**Supplementary Figs. 3b** and **4b**). Taken together, these results represent the first application of amber codon suppression and SPIEDAC labeling in primary neurons, and they encouraged us to further explore the suitability of using this labeling approach in a range of microscopy studies.

**Fig 4.**
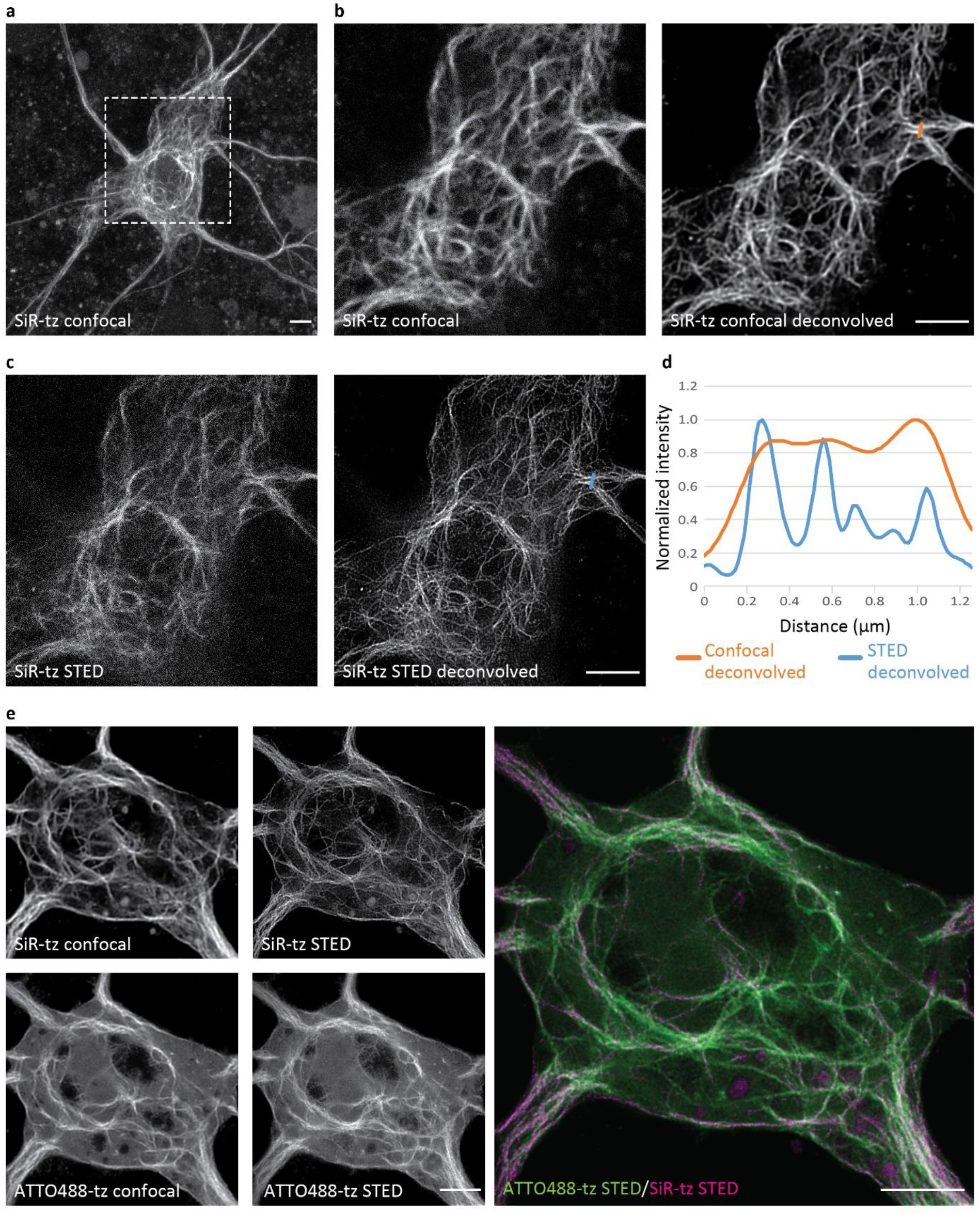
Super-resolution stimulated emission depletion (STED) imaging of click-labeled NFL in primary mouse neurons (MCNs). MCNs expressing NFL^K363TAG^-FLAG, NFM and NES PylRS/tRNA_CUA_^Pyl^ in the presence of TCO‪A-Lys. **a**–**c**, Two to three days after transfection, neurons were labeled with SiR-tz, fixed and stained with anti-FLAG antibody. Afterwards, neurons were imaged with STED super-resolution microscopy. Panel **a** shows a maximum projection of a confocal *Z*-stack of the neuron. The next four images show the confocal (**b**) and STED (**c**) micrographs of the region within the dashed box before and after deconvolution. **d**, The increase in resolution is visualized in a graphical representation of signal intensities across the line profiles drawn in the deconvolved confocal (orange line) and STED (blue line) images. **e**, STED imaging of two populations of click-labeled NFL. Two days after transfection, neurons were labeled with SiR-tz and incubated for a further 2 days with TCO‪A-Lys. Afterwards, cells were fixed, labeled with ATTO488-tz and imaged with STED super-resolution microscopy. Raw confocal and STED images were deconvolved using Huygens deconvolution software. Scale bars: 5 µm (**a**–**c, e**).

### Click labeling allows imaging of NFL in living neurons

Since a number of cell-permeable dyes showed specific fluorescent labeling of NFL^K3636TAG→UAA^ (**Supplementary Figs. 4a–g**), we tested the compatibility of our labeling approach with live-cell imaging. We successfully imaged living neurons expressing NFL^K363TAG→TCO‪A-Lys^ that we labeled with one of the cell-permeable dyes, including tetrazine derivatives of BODIPY, TAMRA, and SiR (**Fig. 2b**). A comparison of labeling quality in live-imaging experiments revealed that different dyes emit different background levels, with SiR- and BODIPY-conjugated tetrazines affording a higher signal-to-noise ratio than TAMRA-tetrazine. In addition, while performing the live-cell imaging, we observed a nonspecific signal in cytoplasmic vesicles, which was not present in images of fixed cells (**Fig. 2b**). Control experiments with SiR-tetrazine revealed that lysosomes accumulated UAA molecules that were labeled with tetrazine-dyes independently of the amber codon suppression (**Supplementary Fig. 5**). This was confirmed by incubating non-transfected neurons with UAA and SiR-tetrazine dye (**Supplementary Fig. 5c**). Although SiR-tetrazine has some residual affinity for lysosomes in the absence of a UAA (**Supplementary Fig. 5b,d**), the majority of nonspecific lysosomal labeling seems to occur due to accumulation of UAA in lysosomes (**Supplementary Fig. 5c**). This type of nonspecific labeling is important to keep in mind for future studies, especially if lysosomal proteins are to be labeled, although interestingly, lysosomes are labeled with lower intensity in neurons that express higher levels of NFL^TAG^ and form more extensive neurofilament networks. This can be seen by comparing lower-expressing TAMRA-tetrazine-labeled with higher-expressing SiR- or BODIPY-tetrazine-labeled neurons (**Fig. 2b**). In addition, this background is less apparent after fixation and cell permeabilization (**Fig. 2a**).

**Fig 5.**
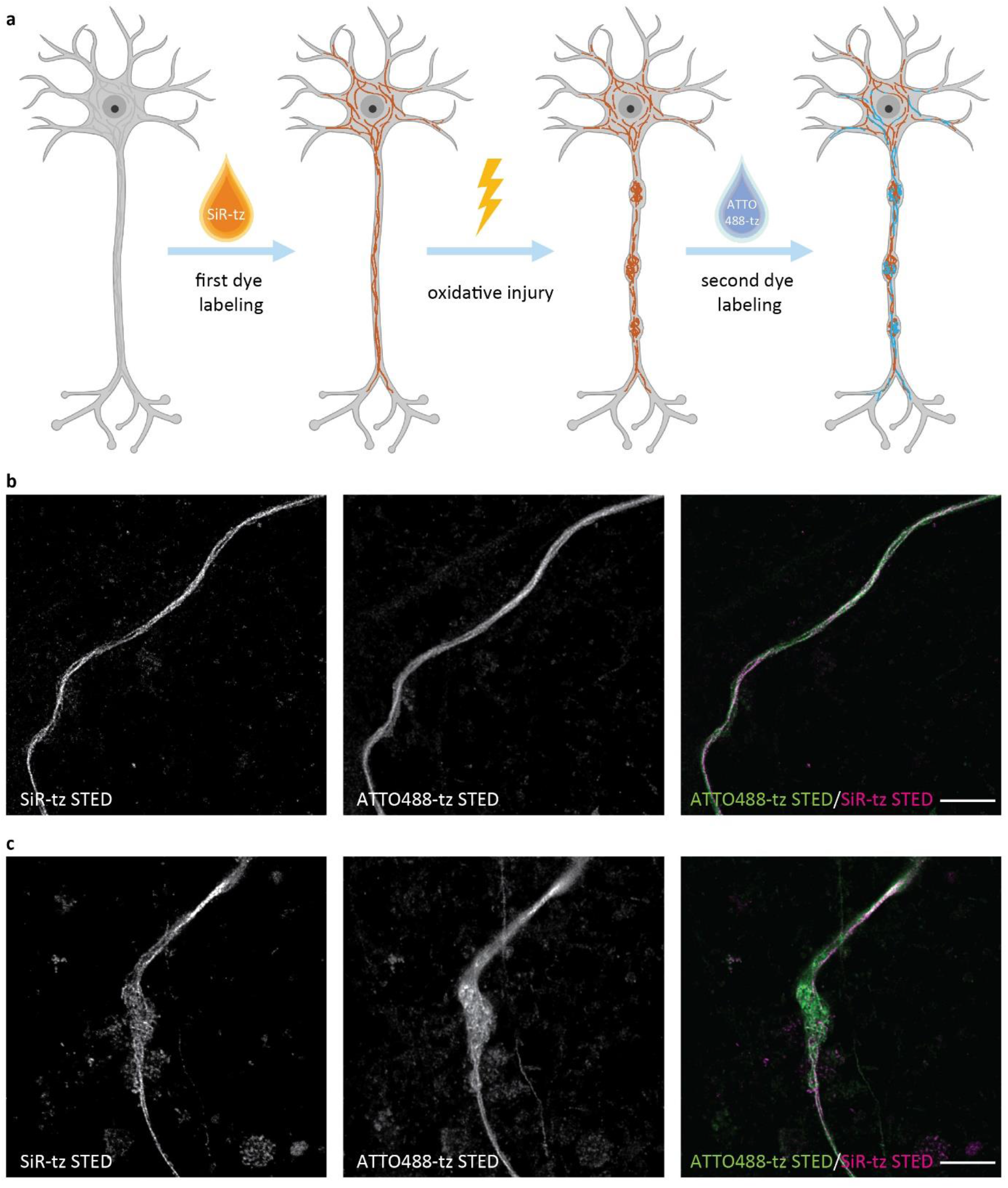
Pulse–chase click labeling of two NFL populations can be used in combination with STED imaging for tracking NFL distribution during oxidative injury. **a**, A schematic representation of the experimental design. MCNs were transfected with NFL^K363TAG^-FLAG, NFM, and NES PylRS/tRNA_CUA_^Pyl^ constructs. After incubation with TCO‪A-Lys for 2 days, NFL was labeled with SiR-tz, and neurons were subjected to nitric-oxide-mediated oxidative injury. After injury, neurons were incubated with TCO‪A-Lys for a further 2 days, fixed, and stained with ATTO488-tz. **b**, STED images of a healthy axon with two populations of click-labeled NFL. **c**, STED images of an injured axon with two populations of click-labeled NFL. STED images were deconvolved using Huygens deconvolution software. Scale bars: 5 µm (**b**,**c**).

### Dual-color pulse–chase click labeling of NFL in primary neurons

One advantage of UAA-based SPIEDAC labeling is the flexibility that it offers with regard to the choice of the tetrazine dye. As shown above, different cell-permeable and cell-impermeable dyes could be used for live-cell and fixed-cell labeling, allowing the possibility of multicolor labeling. We wanted to take advantage of this in order to achieve dual-color labeling in a time-dependent (pulse–chase) manner, as outlined in **Fig. 3a**. By incubating transfected neurons with two tetrazine dyes at two distinct time-points (10 vs. 12 days *in vitro*), we labeled different populations of NFL^K363TAG^, which were synthesized during defined phases of neuronal growth *in vitro*. For these experiments, we combined a pair of cell-permeable dyes. We established a dual-color labeling protocol with a combination of BODIPY-tetrazine and SiR-tetrazine, which allowed us to image two populations of NFL in fixed (**Fig. 3b, Supplementary Fig. 6a**) and living neurons (**Fig. 3c**). For this type of pulse–chase labeling, it is essential that the first dye labels all NFL^TAG^ molecules that have been expressed up to the point the dye is applied. If this happens, then the second dye will label the NFL molecules that were synthesized subsequently. To test this, we performed control experiments in which the second dye showed no labeling when applied immediately after the first dye or after 2 days of incubation in the absence of UAA (**Supplementary Fig. 6b,c**). Although conventional confocal microscopy imaging (**Fig. 3** and **Supplementary Fig. 6**) suggests that most of the neurofilaments in neuronal cell bodies overlap, the potential of this labeling approach lies with studies of target protein trafficking during neuronal growth, injury, or other biological processes.

**Fig 6.**
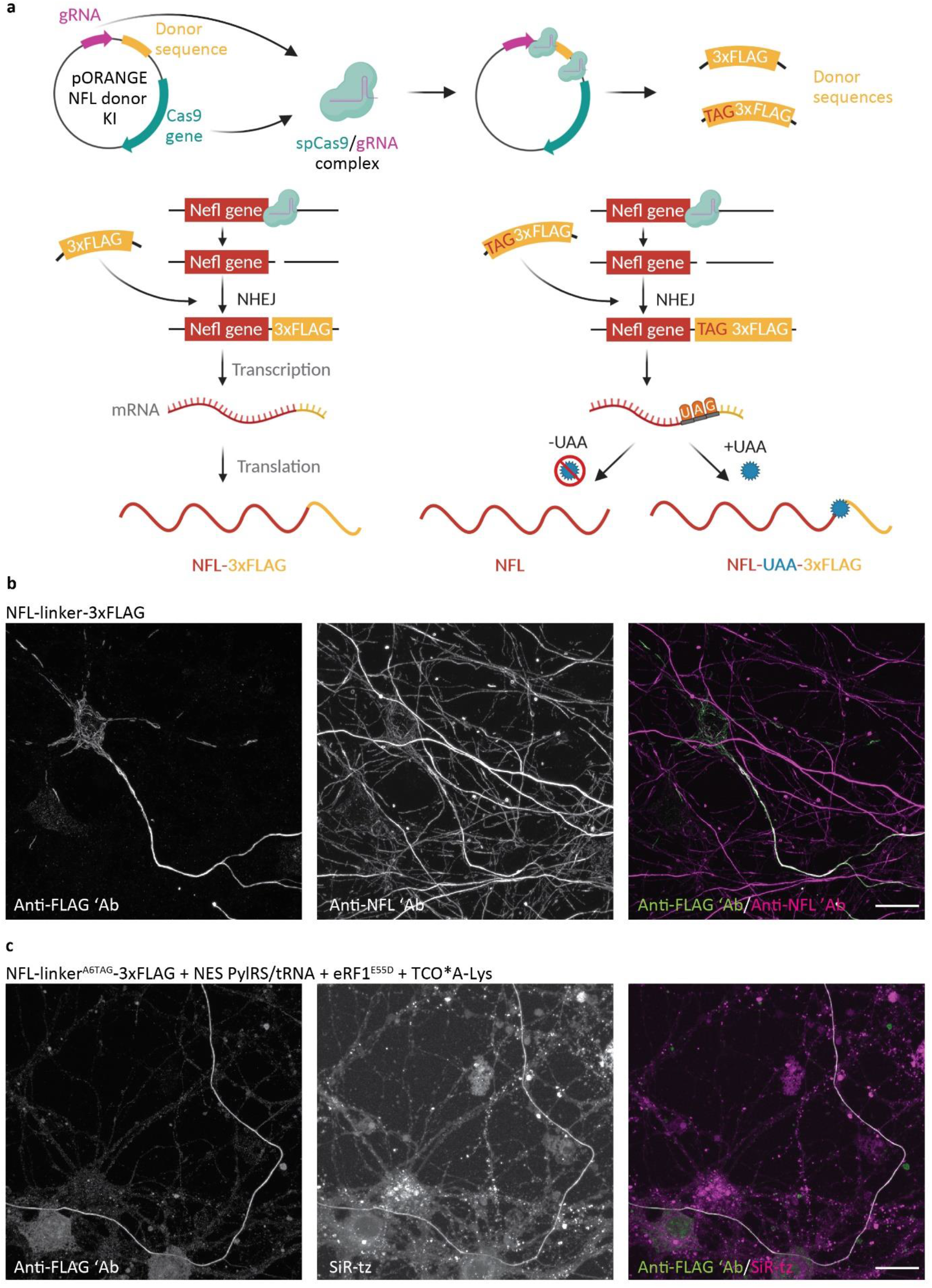
Click labeling of endogenous NFL in primary mouse neurons (MCNs). **a**, A schematic representation of the tagging of endogenous NFL. MCNs were transfected with a pORANGE plasmid encoding for spCas9 and gRNA, and containing linker-3xFLAG or linker^A6TAG^-3xFLAG donor sequence. SpCas9 and gRNA are expressed from the plasmid, cut target sequences around the donor sequence and at the end of the Nefl gene. The resulting donor sequences are used by the non-homologous end joining (NHEJ) system to repair a double strand break in the genomic DNA. Endogenous NFL bearing the linker-3xFLAG sequence is transcribed and translated with the addition of the linker-3xFLAG tag at the C terminus. Endogenous NFL bearing the linker^A6TAG^-3xFLAG tag is fully transcribed. In the absence of a UAA, translation finishes at the UAG codon and full-length NFL is synthesized. In the presence of a UAA and NES PylRS/tRNA_CUA_^Pyl^, the UAA is incorporated in response to the UAG codon, the NFL-linker-UAA-3xFLAG is synthesized, and can be click labeled with a tetrazine dye. **b**, MCNs transfected with pORANGE NFL linker-3xFLAG knock-in construct, stained with anti-FLAG and anti-NFL antibodies, after 6 days of expression. The anti-FLAG and anti-NFL antibodies were stained with AF488- and AF647-conjugated secondary antibodies, respectively. **c**, MCNs transfected with pORANGE NFL linker^A6TAG^-3xFLAG knock-in, NES PylRS/tRNA_CUA_^Pyl^ and eukaryotic release factor 1 mutant E55D (eRF1^E55D^) constructs. After incubation with TCO‪A-Lys for 6 days, the endogenous NFL-linker-UAA-3xFLAG was labeled with SiR-tz, then neurons were fixed and stained with anti-FLAG antibody followed by AF488-conjugated secondary antibody. Scale bars: 20 µm (**b**,**c**).

### Click labeling with minimal fluorescent tags allows super-resolution imaging of NFL in primary neurons

An optimal labeling tag is small and minimizes the impact on the protein of interest. Furthermore, the minimal size of a UAA-based tag, and the possibility to attach the dye directly to the target protein, makes them particularly attractive for super-resolution imaging. To test if SPIEDAC labeling of NFL in primary neurons is compatible with super-resolution imaging, we performed stimulated emission depletion (STED) imaging of SiR-tetrazine-labeled NFL^K3636TAG^. This allowed us to resolve NFL-fibers with improved resolution compared to confocal imaging (**Fig. 4a–d**). In addition, we imaged NFL^K3636TAG^ labeled with ATTO488-tetrazine and CF650-me-tetrazine in neuronal cell bodies (**Supplementary Fig. 7a,b**) and SiR-tetrazine-labeled NFL^K3636TAG^ in neuronal processes (**Supplementary Fig. 7c**). As STED led to improvement in resolution, we combined our pulse–chase dual-color labeling assay with two-color super-resolution imaging. Owing to their compatibility with STED imaging, we used SiR- and ATTO488-tetrazine to resolve two populations of click-labeled NFL in cell bodies (**Fig. 4e**) and neuronal processes (**Supplementary Fig. 8**). Since ATTO488-tetrazine is not cell permeable, we combined live- and fixed-cell SPIEDAC labeling. Finally, to further emphasize the potential of our labeling approach and its suitability for advanced microscopy studies, we combined the dual-color pulse–chase labeling with an oxidative-stress axonal injury (**Fig. 5a**). This allowed us to reveal the distribution of different NFL populations with nanoscale resolution in healthy and injured axons with STED microscopy (**Fig. 5b,c**).

### CRISPR/Cas9 genome editing can be combined with genetic code expansion and SPIEDAC chemistry to achieve labeling of endogenous proteins

To date, click labeling has been achieved by transfecting the POI^TAG^ and expressing it under strong promoters, such as CMV. Although working with transfected cells is routine for cell biologists, overexpression of the POI can affect its function and be toxic for the host cell. We tried to overcome this limitation by combining click labeling and CRISPR/Cas9 genome editing. We used the CRISPR/Cas9-based knock-in strategy based on the recently published ORANGE (Open Resource for the Application of Neuronal Genome Editing) toolbox^30^. As a proof-of-concept, we first mutated the previously described pORANGE vector for tagging endogenous β3-tubulin with GFP. We introduced a Y39TAG mutation in the GFP gene and showed that the full-length β3-tubulin-GFP^Y39TAG^ was only synthesized in the presence of NESPylRS/tRNA^Pyl^ and UAA (**Supplementary Fig. 9**). This demonstrated that amber codon suppression and CRISPR/Cas9 could be combined. We then designed a vector for the C-terminal tagging of endogenous NFL (**Fig. 6a**). Because of the limited efficiency of amber codon suppression, site-specific introduction of TAG at the position K363 would result in knock-down of the endogenous NFL. Therefore, we changed the approach by adding a hexapeptide linker and a 3xFLAG tag at the C terminus of NFL (**Fig. 6a**). In neurons transfected with the construct bearing a linker-3xFLAG donor sequence, tagged endogenous NFL can be detected with anti-FLAG staining (**Fig. 6b**). Co-staining with the anti-NFL antibody confirmed the accuracy of the knock-in, as the anti-FLAG signal completely overlapped with that of anti-NFL (**Fig. 6b**). For the purpose of amber codon suppression and click labeling, we introduced a TAG codon at the position A6 of the linker (linker^A6TAG^; **Fig. 6a**). We co-transfected this construct, together with the NESPylRS/tRNA^Pyl^ and mutant eukaryotic release factor 1 (eRF1 E55D)^31^ to increase the efficiency of amber codon suppression. In the presence of TCO‪A-Lys, the UAG codon is suppressed, and endogenous NFL is tagged with linker-^A6TAG→TCO‪A-Lys^-3xFLAG and can be labeled by click chemistry (**Fig. 6c**). In the absence of TCO‪A-Lys, translation finishes at the UAG codon and the full-length NFL is translated, leaving endogenous NFL levels unaffected. These results show that it is possible to combine genetic code expansion and click labeling with CRISPR/CAas9 genome editing for the labeling of endogenous NFL. The major drawbacks of the current approach are low transfection and amber codon suppression efficiency of the knock-in constructs. We tried to address the latter by co-transfection with eRF1 E55D. Overall, the method could be further improved by using viral vectors instead of conventional transfection methods.

## Discussion

Here, we describe minimal tags for the fluorescent labeling of proteins in living primary mouse neurons. Our labeling approach is based on bioorthogonal SPIEDAC click chemistry between site-specifically incorporated UAAs and tetrazine dyes.

Incorporation of UAAs into target proteins by genetic code expansion allows the introduction of novel chemical and physical properties into biological systems. This emerging protein engineering technique opens a plethora of possibilities for studies of proteins *in vitro* and *in vivo*, at the single-cell and whole-organism levels. UAAs with diverse side chains have been developed: fluorescent UAAs,clickable UAAs with reactive chemical handles for bioorthogonal labeling, photoresponsive UAAs for light-induced control of protein function, and UAAs for the introduction of post-translational modifications, are some examples. Although in its infancy, genetic code expansion has also been applied in neurobiology, as reviewed recently^32^. In 2007, one of the earliest studies on genetic encoding of UAAs in mammalian cells pioneered the now widely used system for the expression of orthogonal tRNAs under the type 3 eukaryotic RNA polymerase III promoters in HEK293T cells and primary neurons^33^. The authors used this approach to site-specifically incorporate *O*-methyltyrosine, an UAA with an extended side-chain, to study the inactivation of voltage-gated potassium Kv1.4 channel in HEK293T cells. As a proof-of-principle, they showed incorporation of *O*-methyltyrosine in a reporter GFP^TAG^ protein in neurons. In the meantime, other types of UAAs were applied in further studies of neuronal proteins, such as irreversible or reversible light-induced control of NMDA- and AMPA-type glutamate receptors with photoresponsive UAAs^34–37^, fluorescent labeling of NMDA receptors^38^ and the Shaker B voltage-dependent potassium channel^39^, as well as the development of reporters for amyloid precursor protein trafficking and processing^40,41^. However, all of these studies were performed using standard cell lines, and the number of studies showing UAA incorporation in neurons has remained limited. In this respect, the fluorescent UAA dansyl-alanine was used in neuronal stem cells to visualize membrane potential^42^. In another study, photocaged cysteine was used to create a light-induced switch for the inwardly-rectifying potassium channel (KiR2.1) in mouse brain slices and living brain^43^. More recent studies reported incorporation of diverse UAAs in a reporter GFP^TAG^ protein with the help of adeno-associated virus (AAV) and baculovirus vectors in dissociated neurons and organotypic slices^44,45^, as well as AAV-based UAA incorporation in mouse brain^44^. We have now expanded the portfolio of genetic code expansion applications in neuroscience by showing that clickable UAAs can be genetically encoded in neurons and used for site-specific fluorescent labeling. As discussed earlier, until now this labeling approach has only been used in standard and readily transfected cell lines and not in complex cells, such as neurons. Here, we show its versatility by performing advanced microscopy studies involving fixed-cell and live-cell imaging, pulse–chase dual-color labeling, and STED imaging of a click-labeled target protein.

We selected the neuron-specific cytoskeletal protein NFL as a target protein. NFL is the smallest of the three major neurofilament subunits. It interacts with neurofilament medium and neurofilament heavy chain, as well as with α-internexin (in the central nervous system) and peripherin (in the peripheral nervous system) to form an extensive network of neurofilaments. Neurofilaments are intermediate filaments with a 10 nm diameter that have important roles in regulating axonal diameter and conduction velocity, and in modulating synaptic plasticity and activity^46,47^. Certain alterations in neurofilament genes, such as Charcot–Marie–Tooth disease-associated NFL mutations, are direct causes of neurological diseases. In other neurological diseases, such as ALS, and giant axon neuropathy, abnormal transport and processing of neurofilaments appear to be a secondary consequence of the disease pathology. Furthermore, NFL levels in the cerebro-spinal fluid and blood correlate with the severity of various neurological conditions, and have been proposed as potential biomarkers of disease progression^48^. Nevertheless, our understanding of neurofilament roles in physiological and pathophysiological conditions remains limited.

Until now, fluorescent labeling of neurofilament subunits has been achieved by making FP fusions. However, fusing an FP to a neurofilament subunit might interfere with neurofilament assembly^49^. Here, we describe an alternative method for labeling NFL in living neurons, which in combination with the existing methods, might spur novel discoveries in the field. Finally, in addition to labeling NFL, our approach offers possibilities for labeling and performing advanced microscopy studies of other neuronal proteins.

To facilitate implementation of click labeling technology for new applications in neurobiology, we provide a workflow for establishing the click labeling of additional target proteins. In this regard, selecting an appropriate host cell line for amber codon suppression is a key prerequisite. As successful amber codon suppression involves creating multiple TAG mutants, their screening and labeling optimization would be too time-consuming in primary neurons. Instead, this can be done in ND7/23 cells, a neuron-like cell line that is transfected with high efficiency. Compared to conventional host cell lines for amber codon suppression, such as HEK293 and COS7, the advantage of ND7/23 cells is that they have neuronal properties^50^, hence they are frequently used as a model for differentiated neurons, for example, in electrophysiological studies. Thus depending on the experimental design, ND7/23 cells can be used either as intermediate hosts for click labeling optimization (as applied in this manuscript) or as a main host for studies involving click-labeled neuronal proteins.

Compared with other methods, SPIEDAC labeling has a number of advantages. Firstly, this is the only method that allows site-specific labeling of proteins with single-residue precision. Clickable UAAs are L-lysine derivatives, adding only a few more atoms to the target protein, thus lowering the risk of functional impact. In addition, SPIEDAC labeling brings the dye as close as possible to the target protein. This is especially relevant for super-resolution imaging^1–5^. Conventional labeling approaches with large antibody complexes can place the fluorophore 20–30 nm away from the target protein. This results in “linkage error”, which introduces localization artefacts and affects localization precision. Although this is not relevant for conventional microscopy with a resolution above 250 nm, it is a problem for super-resolution microscopy, especially if the resolution limit (5-10 nm) reaches the size of conventional labeling tags. UAA-based labeling also allows fluorophores to be placed at a higher density, which is crucial for achieving optimal resolution^51^. Furthermore, our approach is compatible with live-cell imaging as it relies on the rapid and bioorthogonal SPIEDAC click reaction. An additional advantage of this labeling approach is its potential fluorogenic character. Owing to the photophysical properties of certain fluorophores, their tetrazine derivatives are quenched, and their fluorescence is restored/enhanced upon reaction with a click-reactive partner. This property reduces background signal and makes SPIEDAC labeling with tetrazine dyes very attractive for live-cell and super-resolution microscopy applications, as recently reviewed^52^. Moreover, as also shown in our manuscript, SPIEDAC labeling brings flexibility in selecting the tetrazine dye. As different imaging techniques require dyes with different properties—for example, cell-permeable dyes are needed for live-cell imaging, photostable dyes for STED imaging, photoswitchable dyes for single-molecule localization techniques—the most suitable tetrazine dye can be chosen, depending on the desired application. SPIEDAC chemistry can even be adapted for fixed-cell labeling with certain cell-impermeable dyes, albeit at a cost of higher background. In addition, our approach is readily adapted to take advantage of new discoveries in fluorescent dye synthesis^53–56^ and it could be used to attach other types of probes (e.g. gold particles, affinity probes, PET, MRI imaging tracers) to the proteins of interest. Finally, the large variety of fluorophores allows dual-color and potentially multi-color labeling studies. This can be achieved by using different tetrazine dyes, as we described here, or with mutually orthogonal SPIEDAC reactions, as shown previously^24^. In either case, with the addition of UAAs or tetrazine dyes at precisely defined time points, different populations of target proteins can be labeled. With the complex architecture of neurons in mind, we believe that this type of pulse–chase labeling will grant us novel insights into the fates of neuronal proteins.

However, there are also limitations that need to be considered when applying SPIEDAC-based protein labeling. Firstly, amber codon suppression relies on complex genetic engineering machinery that needs to be introduced into the host cell. Depending on the host cell line, this can result in low efficiency of UAA incorporation, especially in cells that are not readily transfected, such as non-dividing neurons. However, as we show with this study, even with conventional transfection methods, we achieved successful incorporation of UAAs in primary neurons. This might differ for other target proteins, but can be further improved by using viral vectors or transgenic animals for the expression of the orthogonal aaRS/tRNA pairs. Another limitation is that our approach relies on amber stop codon suppression, which shows position and sequence context-dependent efficiency^57^. Therefore, different TAG positions need to be tested to find the best-expressing mutant, but also for functional reasons, as site-specific mutations can alter the function of a target protein. However, although this is important for the successful application of SPIEDAC labeling for physiological studies, the site-specificity of the method offers a sufficient chance of finding the most suitable position. In addition, compared to other labeling tags, exchanging one of the residues with a UAA represents a much smaller modification. Another potential limitation is that amber codon suppression might impact the translation of other proteins from genes that naturally use the amber codon as a stop codon. To what extent this happens in mammalian cells is unknown, but as termination of translation depends on additional factors that even compete with incorporation of UAAs at our desired site-specifically introduced TAG site, this type of background should be negligible and is not apparent from microscopy studies. Furthermore, although still at the proof-of-principle stage or with limited applicability in eukaryotic cells, alternative strategies using orthogonal organelles^58^ for amber codon suppression, quadruplet codons^59^ instead of stop codons, and engineered ribosomes^60^, offer solutions to this problem. Another limitation is that our approach and all studies published to date involving SPIEDAC labeling rely on overexpression of the target protein. Whereas this is a standard aspect of many cell biology experiments (not only for SPIEDAC-based labeling, but with other types of genetically encoded probes), protein overexpression can cause artefacts and cytotoxicity. To overcome this, we show here that SPIEDAC chemistry can be combined with CRISPR/Cas9 genome engineering for labeling endogenous neuronal proteins, thus avoiding the unwanted effects of protein overexpression. In this study, we took advantage of the recently described pORANGE vectors for endogenous protein tagging in neurons. To avoid affecting the levels of expression of endogenous NFL, instead of a site-specific TAG knock-in, we introduced the TAG amber codon at the end of the Nefl gene, in the form of a short linker sequence followed by 3x-FLAG. A similar approach was previously used to develop standardized tags for the click labeling of overexpressed proteins^61^. As illustrated in Fig. 6a, our knock-in strategy ensures the level of NFL protein remains unaffected, while allowing SPIEDAC labeling of NFL expressed under the endogenous Nefl-promoter in living neurons. Despite its limited efficiency, this is a first demonstration of fluorescent SPIEDAC labeling of proteins expressed under endogenous promoters. By using more efficient viral vectors, this approach can be further refined and brought beyond the proof-of-principle level. However, it is important to note that when using the pORANGE knock-in strategy for endogenous NFL tagging, the availability of protospacer adjacent motif (PAM) sites was a limiting factor. Since the only suitable PAM site at the end of the Nefl gene is located upstream of the stop codon, our approach resulted in deletion of six C-terminal amino acids. This limitation can be avoided by using novel genome engineering strategies that rely on targeting of noncoding regions, and which offer more flexibility in choosing the target sites^62^.

In summary, we have established a novel approach for live-cell protein labeling in primary neurons by combining two state-of-the-art techniques: incorporation of UAAs via genetic code expansion and ultrafast bioorthogonal SPIEDAC reactions. Site-specific labeling with UAA-based minimal tags expands the toolbox of available live-cell protein labeling methods in neurons. Furthermore, by establishing SPIEDAC labeling in neurons, we further expanded the portfolio of genetic code expansion-based applications in neuroscience. We believe that by complementing the currently available methods, labeling by click chemistry will open new possibilities for advanced studies of neuronal cells involving neuronal protein labeling, trafficking in living neurons, labeling of endogenous proteins, and advanced fixed-cell and live-cell super-resolution studies.

## Materials and methods

### Cell culture

Mouse neuroblastoma x rat neuron hybrid ND7/23 cells were purchased from Sigma-Aldrich (ECACC 92090903). They were grown in high-glucose Dulbecco’s Modified Eagle Medium (DMEM; Thermo Fisher Scientific, cat. no. 41965062) supplemented with 10% heat-inactivated fetal bovine serum (Thermo Fisher Scientific, cat. no. 10270106), 1% penicillin-streptomycin (PS; Sigma Aldrich, cat. no. P0781), 1% sodium pyruvate (Thermo Fisher Scientific, cat. no. 11360039) and 1% L-glutamine (Thermo Fisher Scientific, cat. no. 25030024). FBS was inactivated by incubation at 56 °C for 30 min. Cells were passaged three times per week, and used for transfections at passages 3 to 15.

For microscopy experiments, ND7/23 cells were seeded on eight-well Lab-Tek II chambered coverglasses (German #1.5 borosilicate glass; Thermo Fisher Scientific, cat. no. 155409) at a density of 25,000 cells per well. Prior to cell seeding, the coverglasses were coated with a 10 µg/ml solution of poly-D-lysine (Sigma-Aldrich, cat. no. P6407) in double-distilled water (ddH_2_O) for a minimum of 4 h at room temperature (RT). Chambered coverglasses were washed three times with ddH_2_O and allowed to dry prior to cell seeding.

Primary mouse cortical neurons (MCNs) from C57BL/6 embryonic day 17 were purchased from Thermo Fisher Scientific (cat. no. A15586). They were thawed and cultured according to the manufacturer’s recommendation in a B-27 Plus Neuronal Culture System consisting of Neurobasal Plus (NB Plus) medium and B27 Plus supplement (Thermo Fisher Scientific, cat. no. A3653401). Culturing medium was prepared by adding 2% of B27 Plus supplement and 1% of PS to Neurobasal Plus + (NB Plus +). For widefield and confocal microscopy experiments, MCNs were seeded on eight-well Lab-Tek II chambered coverglasses at a density of 90,000 cells per well. For experiments involving STED imaging, MCNs were seeded on eight-well µ-slides with glass bottoms (Ibidi cat. no. 80827), at a density of 100,000 cells per well. The glass bottoms of the Lab-Tek chambers and µ-slides were pre-coated with a 20 µg/ml solution of poly-D-lysine in ddH_2_O for 2 h at RT. Prior to cell seeding, Lab-Tek coverglasses and µ-slides were washed three times with ddH_2_O, allowed to dry, and the pre-incubated for at least 30 min with NB Plus + medium. During the culturing of the MCNs, half the NB Plus + medium was exchanged twice per week.

### Constructs, cloning, and mutagenesis

The cDNA encoding for mouse neurofilament light chain (NFL) was amplified from the vector pmNFL (a gift from Anthony Brown, Addgene plasmid #83127; http://n2t.net/addgene:83127;RRID:Addgene_83127)^63^ and initially cloned in an mEGFP-N1 plasmid (a gift from Michael Davidson, Addgene plasmid #54767; http://n2t.net/addgene:54767;RRID:Addgene_54767) using HindIII (Thermo Fisher Scientific, cat. no. FD0504) and ApaI (Thermo Fisher Scientific, cat. no. FD1414) enzymes. In the resulting construct, the TAG amber stop codon was introduced at positions K211, K363, R438, and K468 of the NFL cDNA, via PCR-based site-directed mutagenesis. After the mutagenesis, GFP was excised from all constructs using the enzymes BamHI (Thermo Fisher Scientific, cat. no. FD0054) and NotI (Thermo Fisher Scientific, cat. no. FD0595), and replaced by a double-stranded DNA oligonucleotide containing the FLAG tag sequence (DYKDDDDK). The FLAG tag oligonucleotide was synthesized by Sigma-Aldrich as two complementary single-stranded oligonucleotides (Supplementary Table 1).

Together with NFL, we co-transfected neurofilament medium chain (NFM) cDNA-containing plasmid pmNFM (a gift from Anthony Brown, Addgene plasmid #83126; http://n2t.net/addgene:83126;RRID:Addgene_83126)^63^.

For the experiments involving amber codon suppression of overexpressed NFL^TAG^ mutants, we used a previously published pcDNA3.1/Zeo(+) plasmid^25^ containing cDNA that encodes *Methanosarcina mazei* pyrrolysyl tRNA synthetase with a nuclear export signaling sequence and Y306A, Y384F substitutions (NES PylRS^AF^), and one copy of tRNA_CUA_^Pyl^ under the control of the U6 promoter (a kind gift from Edward Lemke’s laboratory, EMBL, Heidelberg, and IMB, Mainz).

For the experiments involving amber codon suppression of endogenous NFL and βIII tubulin, we used a pcDNA3.1/Zeo(+) plasmid containing codon-optimized cDNA that encodes *Methanosarcina mazei* NES PylRS^AF^ and one copy of tRNA_CUA_^Pyl^. The codon-optimized cDNA encoding the NES PylRS was synthesized by GenScript and cloned into the pcDNA3.1/Zeo(+) vector. We subsequently added tRNA_CUA_^Pyl^ and the U6 promoter upstream of the CMV promoter in the reverse direction by cloning, using BglII (Thermo Fisher Scientific, cat. no. FD0083) and MfeI (Thermo Fisher Scientific, cat. no. FD0753) enzymes. The U6 promoter-tRNA cassette synthesized by GenScript was used as a template for the cloning. For the amber codon suppression of endogenous NFL and βIII tubulin, we also co-transfected neurons with eukaryotic release factor 1 E55D mutant (eRF1^E55D31^). This plasmid was cloned by Christopher D. Reinkemeier in Edward Lemke’s laboratory.

For the labeling of endogenous βIII tubulin, we used a pORANGE Tubb3-GFP KI plasmid (a gift from Harold MacGillavry, Addgene plasmid #131497; http://n2t.net/addgene:131497;RRID:Addgene_131497)^30^. For the optimization of genetic code expansion of endogenous βIII tubulin, we replaced GFP from this construct with GFP^Y39TAG^ by cloning with HindIII (Thermo Fisher Scientific, cat. no. FD0504) and XhoI (Thermo Fisher Scientific, cat. no. FD0694) restriction sites.

In order to label endogenous NFL, we designed and cloned target and donor sequences following the previously published protocol^30^. The NFL target sequence GAGTGCTGGAGAGGAGCAGG (https://wge.stemcell.sanger.ac.uk//crispr/377510968) was selected using the Ensembl browser^64^ [Ensembl release 102, November 2020; Mus musculus version 102.38 (GRCm38.p6), Chromosome 14: 68,087,408–68,087,430] based on available PAM sites at the end of the Nefl gene. The integration site is Q537 and the knock-in results in the deletion of six C-terminal amino acids of the NFL protein. The target sequence was subsequently cloned into the pORANGE cloning template vector (a gift from Harold MacGillavry Addgene plasmid #131471; http://n2t.net/addgene:131471;RRID:Addgene_131471)^30^ using the BbsI enzyme (Thermo Fisher Scientific, cat. no. FD1014) and single-stranded DNA oligonucleotides synthesized by Sigma-Aldrich. In the next step, we cloned a donor sequence containing *linker*-3xFLAG (*GSAGSA*-DYKDHDGDYKDHDIDYKDDDDK) or *linker*^*A6TAG*^-3xFLAG (*GSAGS‪-*DYKDHDGDYKDHDIDYKDDDDK) into the resulting plasmid (pORANGE NfL KI), using the enzymes HindIII and BamHI. The donor sequences were amplified by PCR from existing plasmids.

All primer and oligonucleotide sequences used for cloning and mutagenesis are listed in Supplementary Table 1.

### UAAs, tetrazine derivatives of fluorescent dyes, and antibodies

In this study, the following unnatural amino acids (UAAs) were used: *trans*-cyclooct-2-en-L-lysine (TCO‪A-Lys; Sirius Fine Chemicals, SICHEM, cat. no. SC-8008), *trans*-cyclooct-4-en-L-lysine (TCO4en/eq-Lys; a kind gift from Edward Lemke’s laboratory, also available from SICHEM, cat. no. SC-8060) and *endo*-bicyclo[6.1.0]nonyne-lysine (endo-BCN-Lys; SICHEM cat. no. SC-8014). For SPIEDAC labeling, the following tetrazine derivatives of fluorescent dyes were used: ATTO655-methyltetrazine (ATTO655-me-tz; ATTO-TEC GmbH, cat. no. AD 655-2502), ATTO488-tetrazine (ATTO488-tz; Jena Bioscience, cat. no. CLK-010-02), CF500-methyltetrazine (CF500-me-tz; Biotium cat. no. 96029), CF650-methyltetrazine (CF650-me-tz; Biotium cat. no. 96036), Janelia Fluor 646-methyltetrazine (JF646-me-tz; Jena Bioscience custom synthesis), Janelia Fluor 549-tetrazine (JF549-tz; Tocris cat. no. 6502), silicon rhodamine-tetrazine (SiR-tz; SpiroChrome cat. no. SC008), Alexa Fluor 647-tetrazine (AF647-tz; a kind gift from Edward Lemke’s laboratory), TAMRA-tetrazine (TAMRA-tz; Jena Bioscience cat. no. CLK-017-05), and BODIPY-tetrazine (BODIPY-tz; Jena Bioscience cat. no. CLK-036-05). For immunocytochemistry, the following antibodies were used: rabbit anti-FLAG antibody (Merck Millipore cat. no. F7425), mouse anti-neurofilament 70 kDa antibody, clone DA2 (Merck Millipore cat. no. MAB1615, goat anti-rabbit Alexa Fluor (AF) 488 Plus (Thermo Fisher Scientific, cat. no. A32731), goat anti-rabbit AF555 (Thermo Fisher Scientific, cat. no. A21429), goat anti-rabbit AF647 Plus (Thermo Fisher Scientific, cat. no. A32733), and goat anti-mouse AF647 Plus (Thermo Fisher Scientific, cat. no. A32728).

### Transfection

Both ND7/23 cells and MCNs were transfected using the Lipofectamine 2000 transfection reagent (Thermo Fisher Scientific, cat. no. 11668027). ND7/23 cells were transfected 14-20 h after seeding into an eight-well Lab-Tek chambered slide, with a slightly modified manufacturer’s protocol using a DNA/Lipofectamine 2000 ratio of 1 µg:2.4 µl and up to 0.625 µg of total DNA per well. Immediately after transfection, a stock solution of UAA (100 mM in 0.2 M NaOH containing 15% DMSO) was diluted 1:4 in 1 M HEPES (Thermo Fisher Scientific, cat. no. 15630080) and added to cells to a final concentration of 250 µM for TCO‪A-Lys and TCO4en/eq-Lys, and 1 mM for endo-BCN-Lys. The medium was replaced after incubation for 6 h (37 °C, 5% CO_2_), and the HEPES-diluted UAA was again added before cells were incubated overnight (37 °C, 5% CO_2_).

For the transfection of MCNs, we adapted a previously published protocol^65^using Lipofectamine 2000. As was done for ND7/23 cells, we used a DNA/Lipofectamine 2000 ratio of 1 µg:2.4 µl. The total amount of DNA per well in an eight-well Lab-Tek chambered slide was up to 1.25 µg. DNA and Lipofectamine 2000 solutions were prepared in NB Plus medium with 1% of PS. Prior to their addition to the cells, transfection solutions were mixed with an equal volume of warm NB Plus medium containing 4% B27 Plus to obtain a final B27 Plus content of 2%. These final transfection mixtures were then warmed by incubation for 5 min (37 °C, 5% CO_2_). The culturing medium was aspirated from neurons and retained for use as conditioned medium (CM), and warm transfection mixture was added dropwise to cells. After incubation for 5-6 h, the transfection medium was aspirated, and the retained CM was added back to cells. If the transfection was performed on the day of medium change, half the volume of CM was put back to the cells and topped up with fresh NB Plus + medium. Afterwards, 100 mM TCO‪A-Lys stock (in 0.2 M NaOH containing 15% DMSO) was diluted 1:4 in 1 M HEPES and added to neurons, to a final concentration of 250 µM. MCNs were incubated (37 °C, 5% CO_2_) for a minimum of two days prior to click labeling. For NFL^TAG^ overexpression experiments, MCNs were transfected at day *in vitro* (DIV) 8, and for the labeling of endogenous NFL and βIII tubulin with pORANGE vectors, MCNs were transfected at DIV5.

### Single-color click-chemistry labeling in live ND7/23 cells and neurons with cell-permeable dyes

ND7/23 cells expressing NFL^WT^-FLAG or NFL^TAG^-FLAG mutants were labeled by SPIEDAC click chemistry after overnight (18-20 h) incubation with the UAA. To wash out excess UAA, medium containing the UAA was removed, cells were washed twice with culturing medium and then incubated 2 h in fresh culturing medium. Cells were washed once more with culturing medium and incubated for 10 min (37 °C, 5% CO_2_) with the tetrazine dye diluted in culturing medium. The concentration of tetrazine dyes was 5 µM, except for JF549-tz, which was used at a concentration of 2.5 µM. After incubation for 10 min, the dye-containing medium was aspirated, cells were washed twice and then incubated for 2 h in fresh culturing medium. Afterwards, the culturing medium was aspirated, cells were washed once with 0.01 M phosphate-buffered saline (PBS; 137 mM NaCl, 10 mM Na_2_HPO_4_, 1.8 mM KH_2_PO_4_, 2.7 mM KCl, pH 7.4) and fixed with 4% paraformaldehyde (PFA; Sigma–Aldrich, cat. no. 158127) in 0.1 M phosphate buffer (PB) for 15 min at RT. After fixation, the FLAG tag was labeled by immunocytochemistry.

Live-cell click-chemistry labeling of MCNs was performed 2-3 days (overexpression experiments) or 6 days (labeling of endogenous NFL with pORANGE vectors) after transfection. First, medium containing the TCO‪A-Lys was removed, neurons were washed twice with fresh NB Plus +, and then incubated for 2–3 h (37 °C, 5 % CO_2_) in a 1:1 mixture of fresh NB Plus + and CM (collected either on the day of transfection, or from neurons cultured only for this purpose). Afterwards, neurons were washed once more with fresh NB Plus + and incubated with the tetrazine dye diluted in fresh NB Plus + for 10 min (37 °C, 5% CO_2_). The concentrations of dyes were the same as for the labeling in ND7/23 cells. After the labeling period, neurons were washed twice with fresh NB Plus + and incubated for 2-3h in a 1:1 mixture of fresh NB Plus + and CM. The culturing medium was then aspirated, and neurons were fixed with 4% electron microscopy grade PFA (Electron Microscopy Sciences, cat. no. 15710) diluted in PEM buffer (80 mM PIPES, 2 mM MgCl_2_, 5 mM EGTA, pH 6.8). After fixation, depending on the experiment, and as described in the corresponding figure legends, immunocytochemistry was performed.

### Single-color click-chemistry labeling in ND7/23 cells and neurons after fixation

Labeling with the cell-impermeable tetrazine dyes ATTO655-me-tz, ATTO488-tz, and AF647-tz was performed after cell fixation. First, the UAA was removed and cells were washed according to the washing procedure used for live-cell labeling. After 2-4 h of washing, the medium was aspirated, and ND7/23 cells were rinsed with PBS and fixed with 4% PFA in 0.1 M PB, whereas neurons were fixed without PBS rinsing with 4% PFA in PEM buffer for 15 min at RT. Cells were permeabilized with 0.1% Triton X-100 (Sigma-Aldrich, cat. no. X100) in PBS for 10 min at RT. Tetrazine dyes were diluted to a working concentration of 0.5-2.5 µM in PBS. Cells were rinsed with PBS and labeled with dyes for 10 min at 37 °C. After labeling, cells were rinsed three times with PBS and incubated on a shaker at RT for 20-30 min. Afterwards, the FLAG tag was labeled by immunocytochemistry.

### Pulse-chase click-chemistry labeling of two NFL populations in neurons

Neurons were labeled with the first tetrazine dye (BODIPY-tz) two days after transfection, following the same protocol as for the single-color live click-chemistry labeling. After labeling with the first dye, neurons were washed for 2-3 h in a 1:1 mixture of fresh NB Plus + and CM, and then incubated with TCO‪A-Lys for two further days. After that time, neurons were labeled with the second dye (SiR-tz) by following the same protocol. For the controls of two NFL population labeling, after labeling with the first dye (BODIPY-tz), neurons were either labeled immediately with the second dye (SiR-tz), or incubated without TCO‪A-Lys for 2 days, and then labeled with the second dye (SiR-tz). After labeling with the second dye, neurons were either fixed and stained using anti-FLAG immunocytochemistry or imaged live with confocal scanning microscopy.

For STED imaging experiments involving labeling of two NFL populations and oxidative injury, we established a slightly different protocol. Two days after transfection, neurons were labeled with SiR-tz by following the same protocol as for the single-color live click-chemistry labeling. Then, neurons were washed for 3 h and incubated for 30 min (37 °C, 5% CO_2_) with either 25 µM spermine-NONOate (a nitric oxide donor; Cayman Chemical, cat. no. 82150) or 25 µM sulpho-NONOate (control compound; Cayman Chemical, cat. no. 83300), in the presence of TCO‪A-Lys. After injury, neurons were rinsed with warm NB Plus + and incubated for 2 days with TCO‪A-Lys. Then, neurons were fixed and labeled with 1-1.5 µM ATTO488-tz following the fixed-cell labeling protocol described above.

### MitoTracker and LysoTracker labeling

For the experiments with MitoTracker and LysoTracker labeling, neurons were transfected and labeled with SiR-tz as described above. After washing for 2 h, 250 µl of NB Plus + medium containing 100 nM MitoTracker Orange (Thermo Fisher Scientific, cat. no. M7510) and 400 nM LysoTracker Green (Cell Signaling Technology, cat. no. 8783S) was added to the wells that already contained 250 µl of medium, for final concentrations of 50 and 200 nM for MitoTracker and LysoTracker, respectively. Neurons were incubated with these dyes for 30 min and rinsed twice with NB Plus +. Immediately afterwards, NB Plus + was replaced by Hibernate E medium (Brain Bits LLC, cat. no. HELF) containing 1% PS and 2% B27 Plus, and neurons were imaged live with confocal scanning microscopy.

### Immunocytochemistry staining

For anti-NFL and anti-FLAG immunocytochemical staining, cells and neurons were fixed as described above, then washed three times (5 min each wash) with PBS. Afterwards, cells were incubated with a blocking serum containing 3% bovine serum albumin (Sigma-Aldrich, cat. no. A9647), 10% goat serum (Thermo Fisher Scientific, cat. no. 16210072) and 0.2% Triton X-100 for 1 h at RT. Primary and secondary antibodies were diluted in the blocking serum. Rabbit anti-FLAG antibody was used at a dilution of either 1:1000 (overexpression experiments) or 1:2000 (endogenous NFL labeling experiments with pORANGE vectors). Mouse anti-NFL antibody and all secondary antibodies were used at a dilution of 1:500. Cells were incubated with primary antibodies either overnight at 4 °C or for 1 h at RT, then washed three times (5 min each wash) with PBS and incubated for 1 h at RT with the secondary antibodies. Afterwards, cells were washed three times (5 min each wash) with PBS and either imaged immediately, or stored at 4 °C until imaging.

### Widefield imaging

Widefield epifluorescence imaging was performed on an inverted Nikon Eclipse Ti2-E microscope (Nikon Instruments), equipped with XY-motorized stage, Perfect Focus System, and an oil-immersion objective (Apo 60×, NA 1.4, oil). Setup was controlled by NIS-Elements AR software (Nikon Instruments). Fluorescent light was filtered through 488 (AHF; EX 482/18; DM R488; BA 525/45), 561 (AHF; EX 561/14; DM R561; BA 609/54), and Cy5 (AHF; EX 628/40; DM660; BA 692/40) filter cubes. Afluorescent lamp (Lumencor Sola SE II) was used as a light source and emitted light was imaged with ORCA-Flash 4.0 sCMOS camera (Hamamatsu Photonics). Images were acquired at 16-bit depth, 1024×1024 pixels, and pixel size 0.27 µm.

### Confocal imaging of fixed and live cells

Confocal imaging was performed on a LSM 710 confocal scanning microscope (Zeiss, Oberkochen, Germany) equipped with a Plan-Apochromat 63× objective (NA 1.4, oil), 488, 561 and 633 nm laser lines, and continuous spectral detection. Images were acquired at 16-bit depth, 1024×1024 pixels, pixel size 0.132 µm, with 2× line averaging and a pixel dwell time of 6.3 µs, either as a single plane or as a *Z*-stack with 0.37 µm step size. In all channels, pinhole was set to 1 Airy unit. Emission light was collected sequentially, according to the emission spectra of the fluorophores used.

For the live-cell imaging, a temperature module was used and cells were placed in a heating insert (PeCon, Erbach Germany), which had been equilibrated to 37 °C. The medium used for imaging was Hibernate E medium containing 1% PS and 2% B27 Plus.

### STED imaging

Super-resolution STED imaging was performed on a Leica TCS SP8 microscope (Leica Microsystems IR GmbH, Germany), using an HC PL APO CS2 100×/1.4 oil objective, hybrid detectors and the following laser lines: 488 and 635 nm pulsed excitation lasers, as well as continuous 592 nm and gated pulsed 775 nm depletion lasers. Excitation laser power and detector gain were adjusted for each image individually by avoiding over-saturation, according to the high and low pixel values. Emission light was collected sequentially, according to the emission spectra of the fluorophores used, for example, 500-580 nm (for ATTO488, AF488) or 645-760 nm (for SiR, CF650). Depletion lasers were used at 35-45% of maximum power. Images were acquired at 8-bit depth, image size 2048×2048 pixels, and pixel size 13-14 nm. For the confocal imaging, line averaging was 3× and frame accumulation 3×, whereas for STED imaging frame averaging was from 2-4×, and line accumulation 4×.

### Image processing

Raw images were processed in Fiji software^66^. Widefield images were processed by linear adjustment of brightness and contrast and saved as TIFF files. Confocal *Z-*stack images were converted into maximum intensity projections, the brightness and contrast were adjusted linearly, and images were saved as TIFF files.

STED images and the corresponding confocal images were deconvolved using Huygens deconvolution software (SVI, Netherlands), using Classical Maximum Likelihood Estimation (CMLE) algorithm. The signal-to-noise ratio was set to 20 for confocal and to 7 for STED images, and the maximum number of iterations was 40. Background levels (Supplementary Table 2) were chosen by manually checking the background of each image. After deconvolution, images were imported in Fiji, adjusted linearly for brightness and contrast, and saved as TIFF files.

For presentation purposes, all images were converted to 8-bit depth using Fiji and arranged into figures using Adobe Illustrator. The schemes presented in the paper were made using the BioRender app (BioRender.com) and Adobe Illustrator.

## Acknowledgements

We would like to thank Katja Widmaier for excellent technical assistance, Dr. Adrian Neal for his editorial input, Dr. Edward Lemke and his team for the gift of AF647-tetrazine and plasmids containing NESPylRS^AF^/tRNA pair and eRF1^E55D^, as well as the laboratories of Dr. Anthony Brown, Dr. Michael Davidson and Dr. Harold MacGillavry for sharing additional plasmids which were obtained through Addgene, as described in the Online Methods. We are also grateful to Dr. Harold MacGillavry and his team for helpful discussions about the ORANGE knock-in strategy. This study was supported by the Emmy Noether Programme (project number 317530061 to I.N.-S.) of the German Research Foundation (DFG) and the Werner Reichardt Centre for Integrative Neuroscience (Ministry of Science Baden-Württemberg and former Excellence Cluster EXC307 from the DFG). The Leica STED microscope was funded by a grant from the DFG (INST 2388/62-1).

## Author contributions

A.A. designed and performed the experiments involving ND7/23 cells, primary neurons, widefield and confocal imaging. C.H. performed preliminary experiments involving NFL transfections, mutagenesis and click labeling in ND7/23 cells. N.S. performed preliminary experiments involving genetic code expansion in ND7/23 cells. A.A. and I.N.-S. performed STED imaging, with help from T.S.

A.A. and I.N.-S. analyzed the data, prepared the figures and wrote the manuscript. I.N.-S. conceived and supervised the project. All authors reviewed and approved the final manuscript.

