## Supplementary Information for "Minimal genetically encoded tags for fluorescent protein labeling in living neurons"

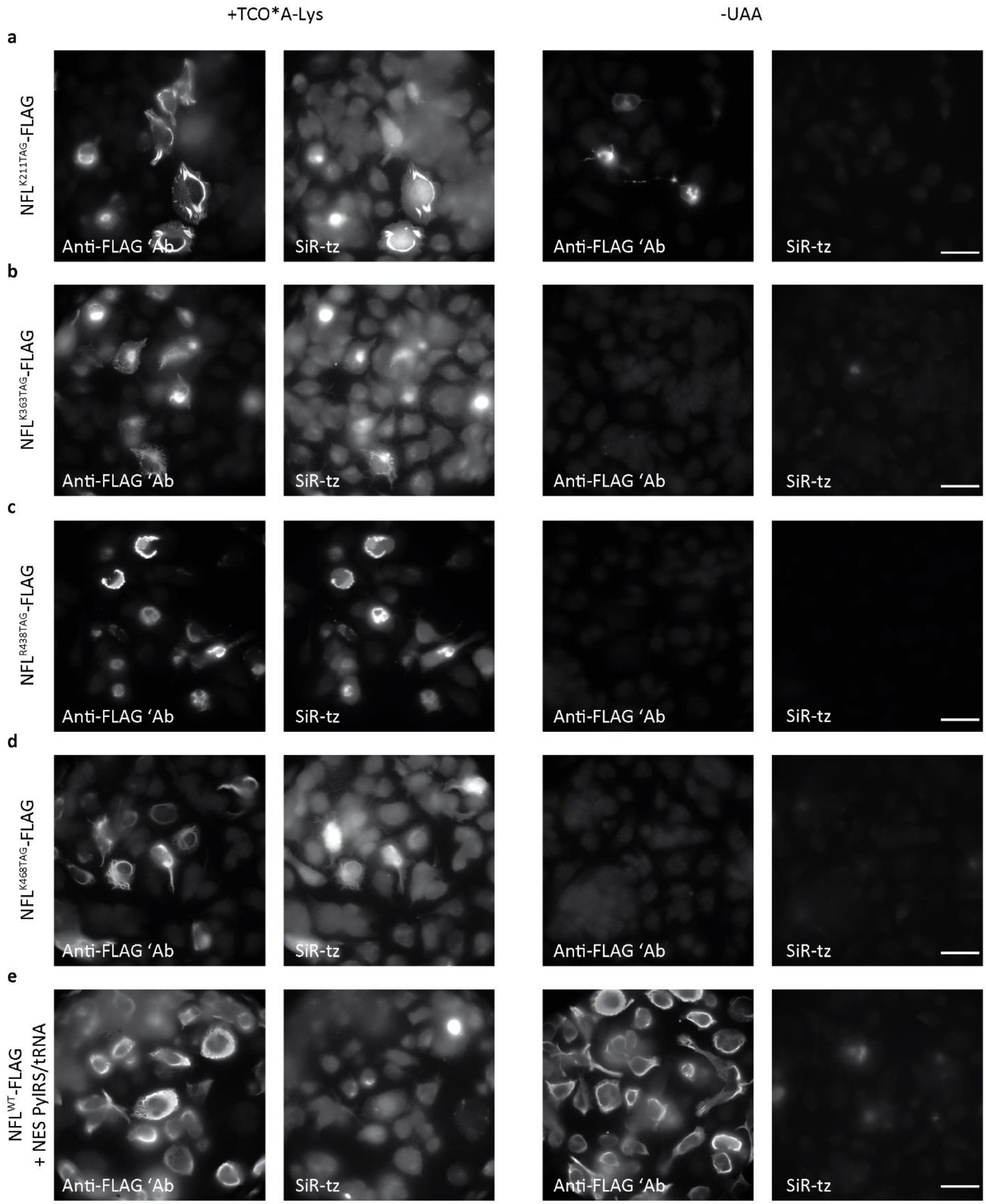


**Supplementary Fig. 1 | Selection of neurofilament light chain (NFL) TAG mutants in ND7/23 cells.** ND7/23 cells expressing NES PylRS/tRNA$_{CUA}^{Pyl}$, NFM, and NFL$^{K211TAG}$-FLAG (**a**), NFL$^{K363TAG}$-FLAG (**b**), NFL$^{R438TAG}$-FLAG (**c**), NFL$^{K468TAG}$-FLAG (**d**) or NFL$^{WT}$-FLAG (**e**). After incubation overnight with or without TCO*A-Lys, cells were labeled with SiR-tetrazine (SiR-tz), fixed and stained with anti-FLAG antibody, followed by Alexa Fluor (AF) 488-conjugated secondary antibody. Images were acquired with widefield microscopy. Scale bars: 50 µm (**a**–**e**).

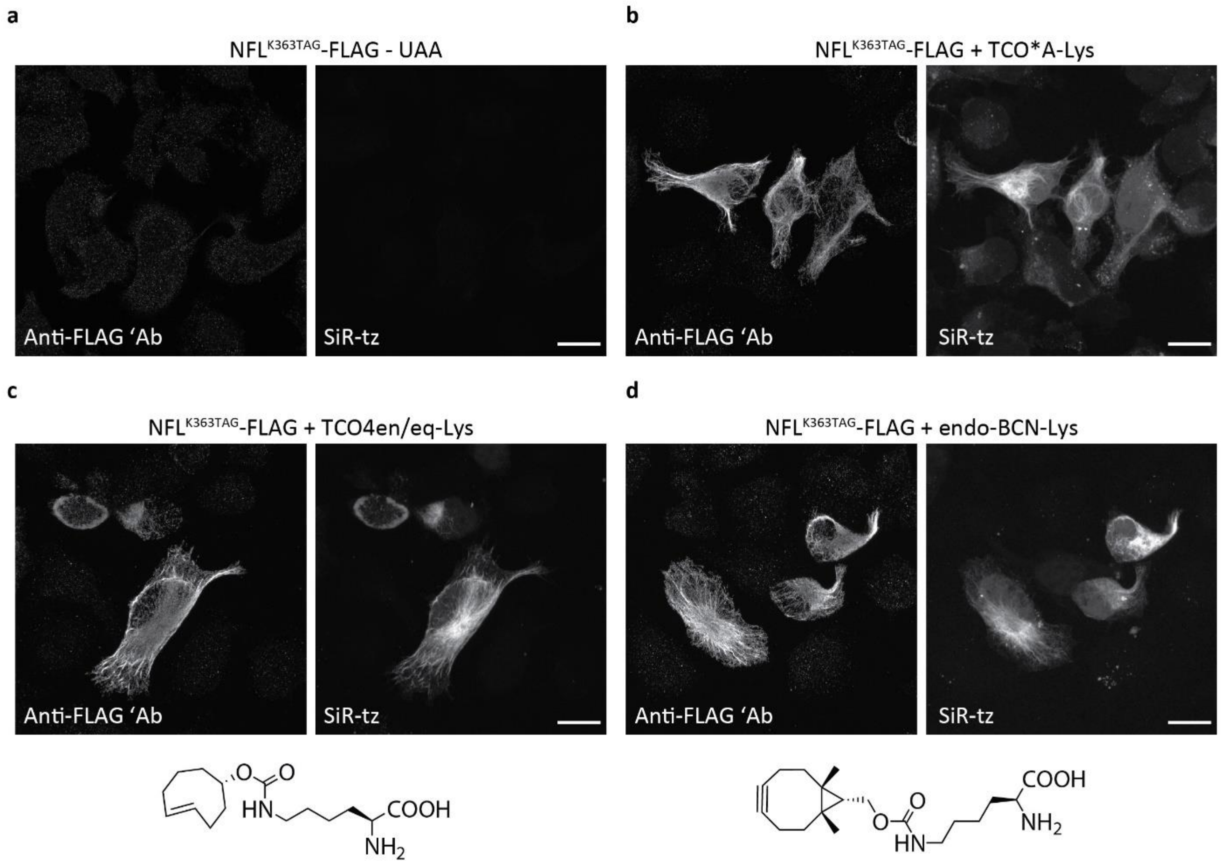


**Supplementary Fig. 2 | Expression of the selected NFL$^{K363TAG}$-FLAG mutant with different unnatural amino acids (UAAs).** ND7/23 cells were transfected with NFL$^{K363TAG}$-FLAG, NFM, and NES PylRS/tRNA$_{CUA}^{Pyl}$ constructs and incubated overnight, either without UAA (**a**), or with TCO*A-Lys (**b**), TCO4en/eq-Lys (**c**), endo-BCN-Lys (**d**). Chemical structures of TCO4en/eq-Lys and endo-BCN-Lys are shown below the corresponding panels. Chemical structure of TCO*A-Lys is shown in the Fig. 1. Cells were labeled with SiR-tz, fixed and stained with anti-FLAG antibody, followed by AF488-conjugated secondary antibody. Images were acquired as a single plane (**a**) or as *Z*-stacks on a confocal scanning microscope (**b**–**d**). *Z*-stack images are shown as maximum intensity projections (**b**–**d**). Scale bars: 20 µm (**a**–**d**).

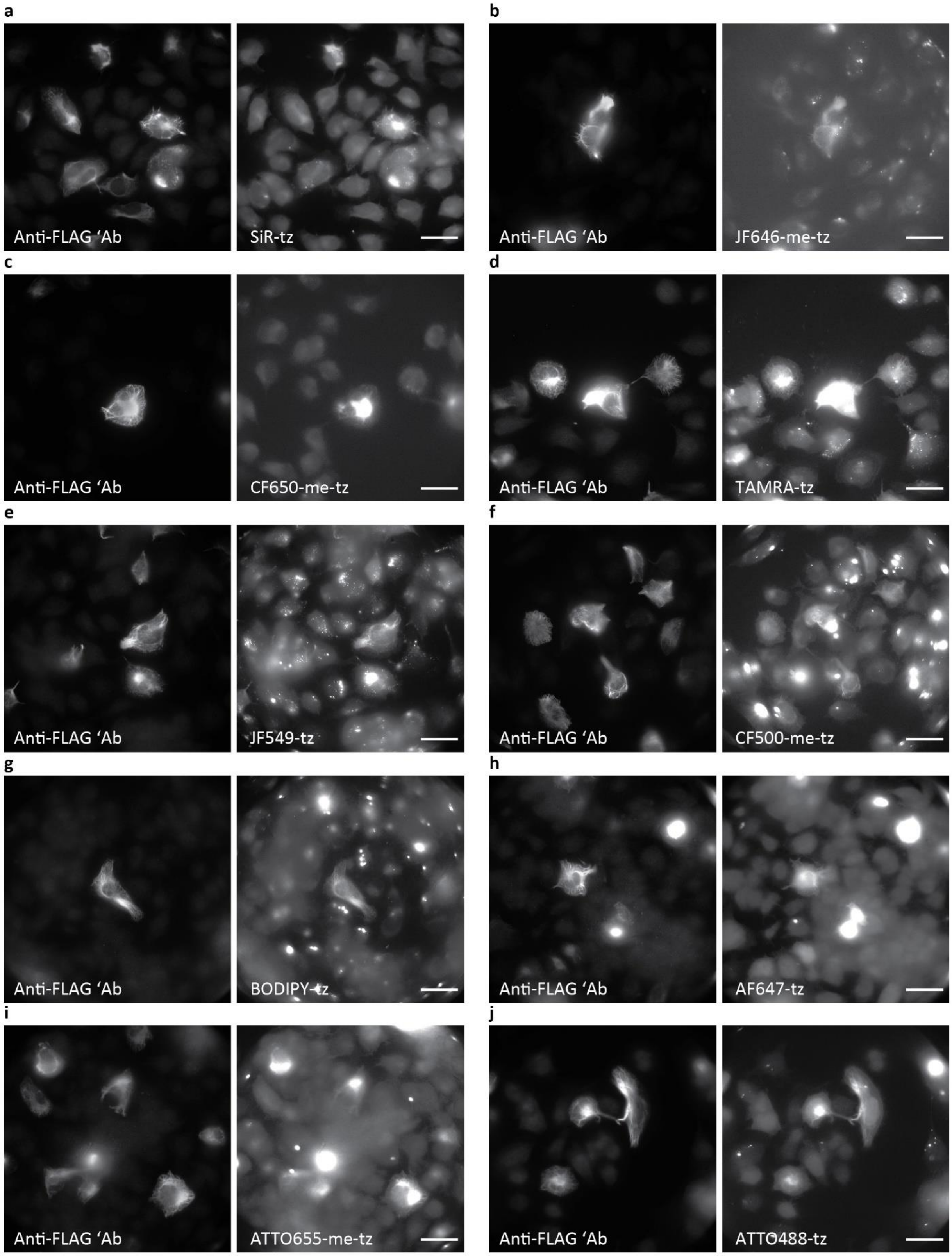


**Supplementary Fig. 3 | Labeling of NFL$^{K363TAG}$-FLAG mutant with various cell-permeable and cell-impermeable tetrazine dyes in ND7/23 cells.** ND7/23 cells expressing NFL$^{K363TAG}$-FLAG, NFM, and NES

PylRS $tRNA_{CUA}^{Pyl}$ constructs. **a**–**g**, After incubation overnight with TCO*A-Lys, living cells were labeled with SiR-tz (**a**), JF646-me-tz (**b**), CF650-me-tz (**c**), TAMRA-tz (**d**), JF549-tz (**e**), CF500-me-tz (**f**), or BODIPY-tz (**g**), then fixed and stained with anti-FLAG antibody. **h**–**j**, After incubation overnight with TCO*A-Lys, cells were fixed and stained with AF647-tz (**h**), ATTO655-me-tz (**i**) or ATTO488-tz (**j**), and with anti-FLAG antibody. Secondary antibodies used for FLAG labeling were conjugated with AF488 (**a**–**e**, **h** and **i**) or AF647 (**f**,**g**,**j**). Images were acquired with widefield microscopy. Scale bars: 50 µm (**a**–**j**).

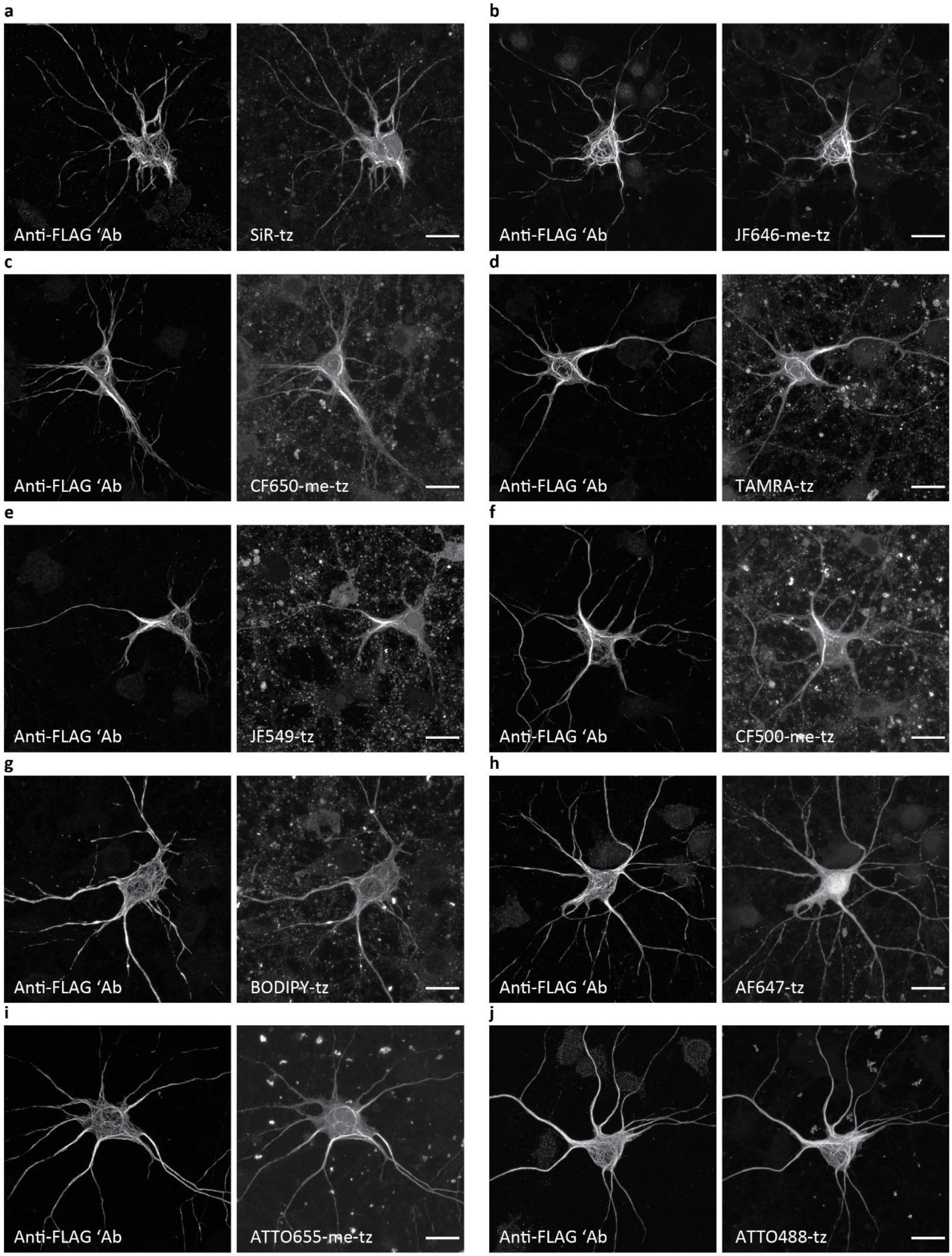


**Supplementary Fig. 4 | Labeling of NFL$^{K363TAG}$-FLAG mutant with various cell-permeable and cell-impermeable tetrazine dyes in primary mouse cortical neurons (MCNs).** MCNs expressing NFL$^{K363TAG}$-

FLAG, NFM, and NES PylRS $tRNA_{CUA}^{Pyl}$ constructs. **a**–**g**, After incubation for 2–3 days in the presence of TCO*A-Lys, living neurons were labeled with SiR-tz (**a**), JF646-me-tz (**b**), CF650-me-tz (**c**), TAMRA-tz (**d**), JF549-tz (**e**), CF500-me-tz (**f**), or BODIPY-tz (**g**), then fixed and stained with anti-FLAG antibody. **h**–**j**, After incubation for 2–3 days in the presence of TCO*A-Lys, neurons were fixed and stained with AF647-tz (**h**), ATTO655-me-tz (**i**) or ATTO488-tz (**j**), and with anti-FLAG antibody. Secondary antibodies used for FLAG labeling were conjugated with AF488 (**a-e**, **h** and **i**), AF647 (**f**,**g**) or AF555 (**j**). *Z*-stack images were acquired on a confocal scanning microscope and are shown as maximum intensity projections. Scale bars: 20 µm (**a**–**j**).

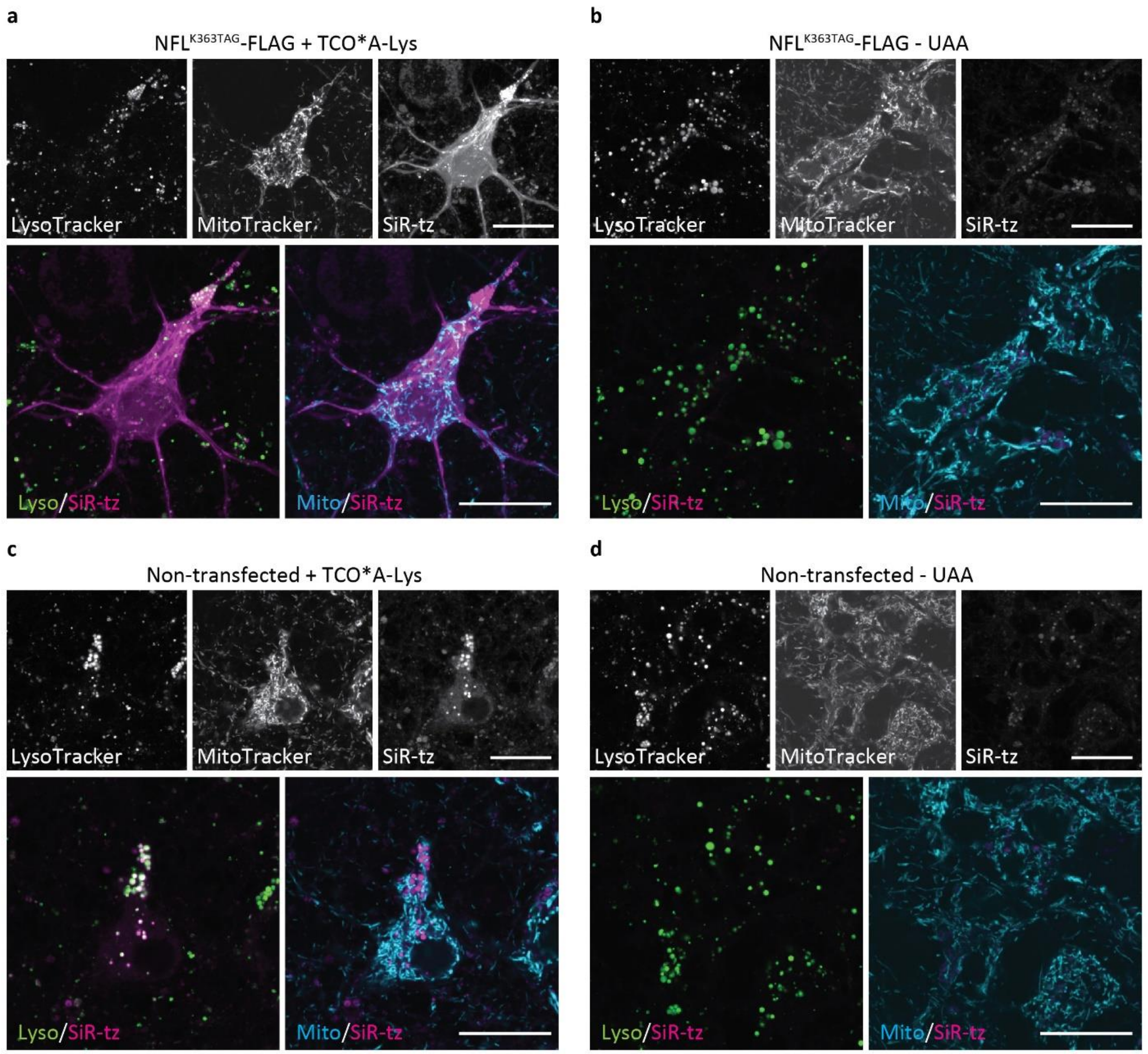


**Supplementary Fig. 5 | Accumulation of tetrazine dyes in lysosomes in the presence of UAA.** Primary mouse cortical neurons were transfected with $NFL^{K363TAG}$-FLAG, NFM, and NES PylRS/$tRNA_{CUA}^{Pyl}$ constructs (**a**,**b**) or with no DNA-containing sham (**c**,**d**). After 3 days of incubation with (**a**,**c**) or without (**b**,**d**) TCO*A-Lys, neurons were labeled with SiR-tz, MitoTracker Orange and LysoTracker Green, and imaged live on a confocal scanning microscope. Brightness and contrast were linearly adjusted to show the same display range in SiR-tz panels, while other panels were adjusted individually for optimal visualization. Scale bars: 20 µm (**a**–**d**).

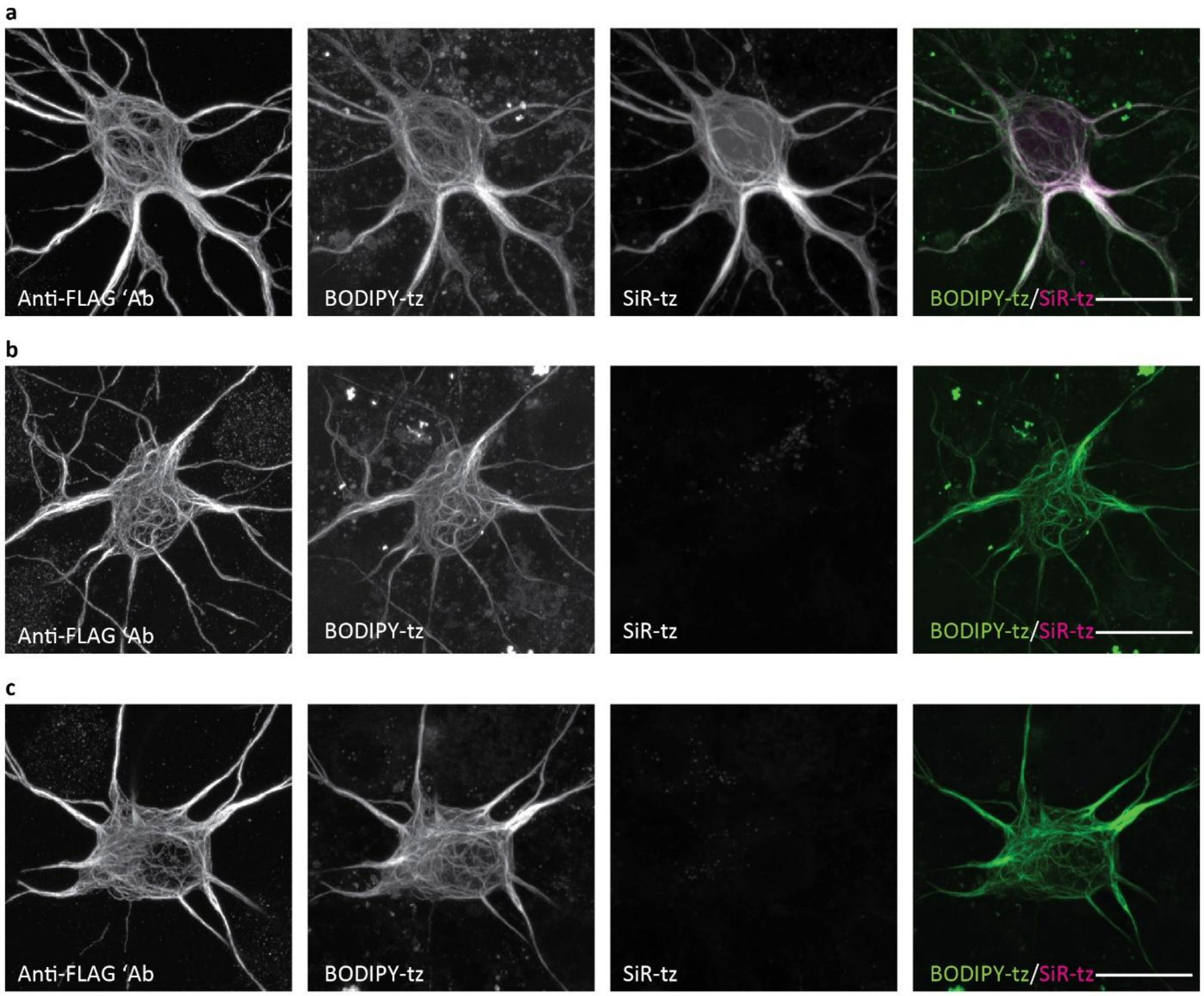


**Supplementary Fig. 6 | Controls for pulse-chase click labeling of two NFL populations in primary mouse cortical neurons (MCNs). a**–**d**, MCNs were transfected with $NFL^{K363TAG}$-FLAG, NFM, and NES PylRS/$tRNA_{CUA}^{Pyl}$ constructs, incubated for 2 days with TCO*A-Lys and labeled with BODIPY-tz dye. After the labeling, neurons were incubated for a further 2 days with TCO*A-Lys (**a**), or without TCO*A-Lys (**b**) and labeled with SiR-tz dye. **c**, Neurons were labeled with SiR-tz immediately after BODIPY-tz labeling. After the second labeling, neurons were fixed and stained with anti-FLAG antibody, followed by AF555-conjugated secondary antibody. *Z*-stack images were acquired on a confocal scanning microscope and are shown as maximum intensity projections. Scale bars: 20 µm (**a**–**c**).

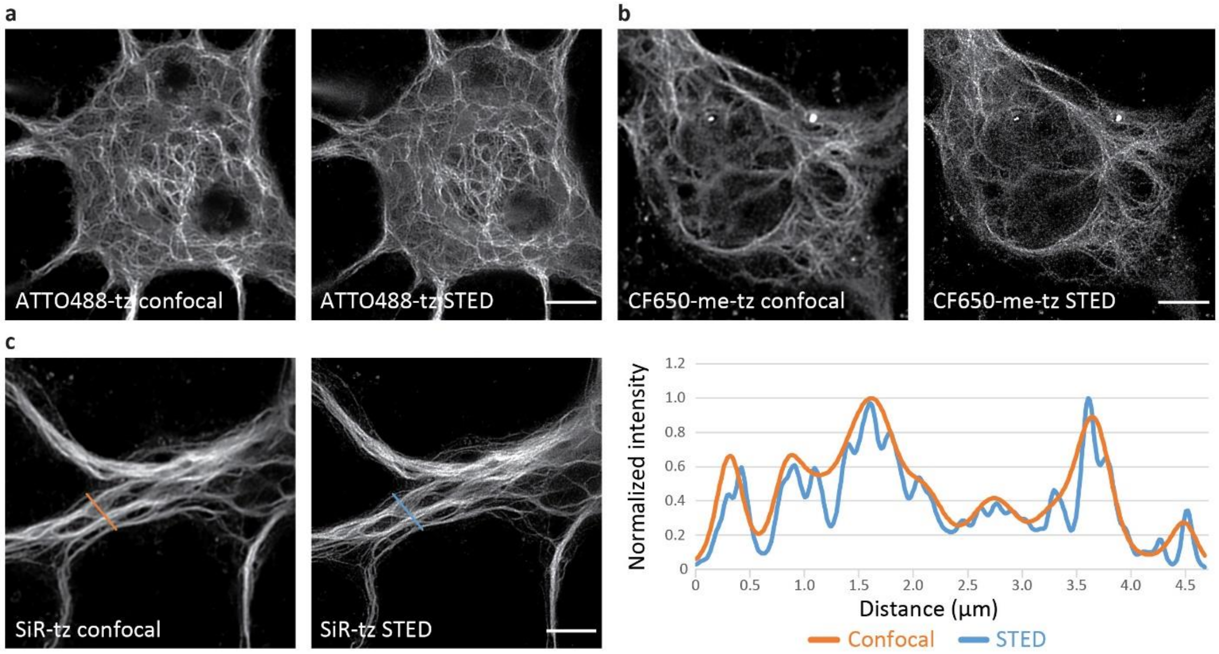


**Supplementary Fig. 7 | Additional examples of stimulated emission depletion (STED) imaging of NFL in primary mouse cortical neurons (MCNs). a**–**c**, MCNs expressing $NFL^{K363TAG}$-FLAG, NFM, and NES PylRS/$tRNA_{CUA}^{Pyl}$ in the presence of TCO*A-Lys. After 2–3 days of expression, NFL was labeled with ATTO488-tz (**a**), CF650-me-tz (**b**) or SiR-tz (**c**) and imaged with STED microscopy. The line profile graph in panel **c** demonstrates the increase in resolution of STED image (blue line) in comparison to the confocal image (orange line). Raw confocal and STED images were deconvolved using Huygens deconvolution software. Scale bars: 5 µm (**a**–**c**).

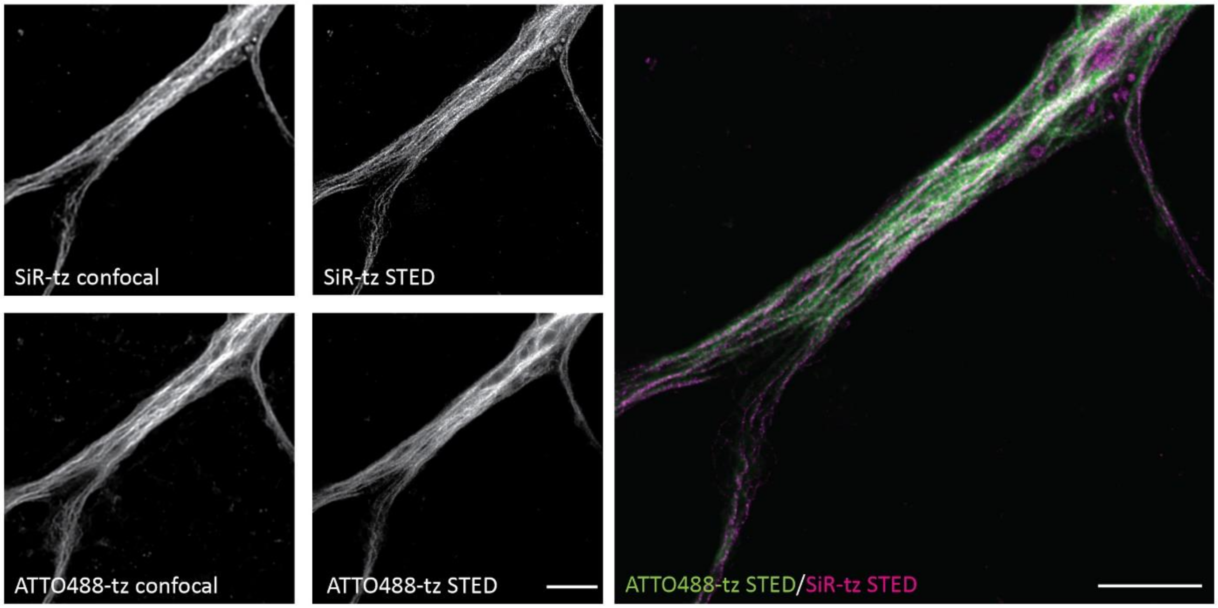


**Supplementary Fig. 8 | Additional examples of STED imaging of two populations of click-labeled NFL in primary mouse cortical neurons (MCNs).** MCNs expressing $NFL^{K363TAG}$-FLAG, NFM, and NES $PylRS/tRNA_{CUA}^{Pyl}$ in the presence of TCO*A-Lys. After 2 days of expression, neurons were labeled with SiR-tz, incubated for a further two days with TCO*A-Lys, and labeled after fixation with ATTO488-tz. Click-labeled NFL was imaged with STED microscopy. Raw confocal and STED images were deconvolved using Huygens deconvolution software. Scale bars: 5 µm.

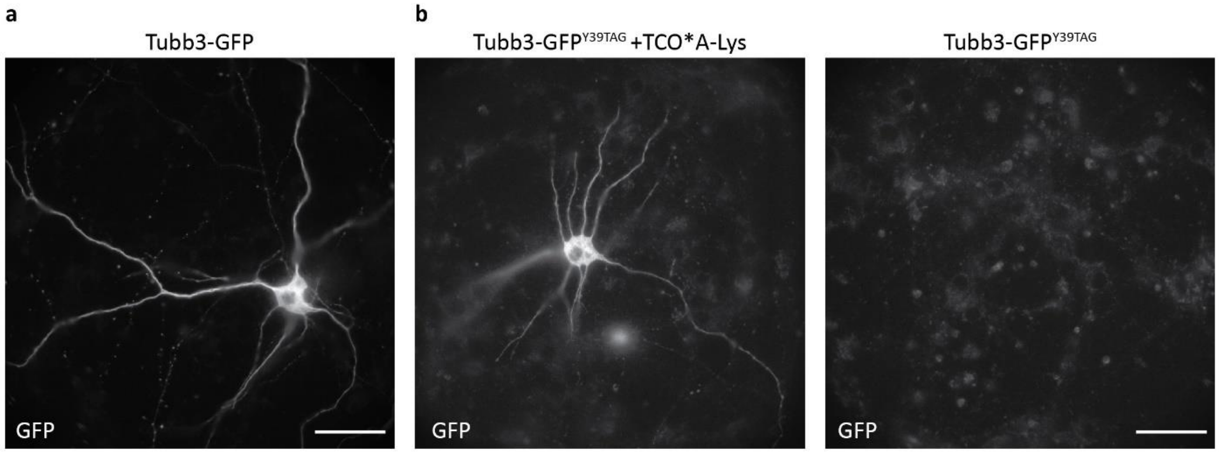


**Supplementary Fig. 9 | Optimization of amber codon suppression of endogenous βIII tubulin (Tubb3) in primary mouse cortical neurons (MCNs). a**, MCNs were transfected with the pORANGE Tubb3-GFP knock-in construct, together with the NES PylRS/tRNA$_{CUA}^{Pyl}$ and eukaryotic release factor 1 mutant E55D (eRF1$^{E55D}$) at day *in vitro* 3. After 6 days of expression, neurons were fixed and endogenous Tubb3 tagged with GFP was imaged with widefield microscopy. **b**, MCNs were transfected with the pORANGE Tubb3 GFP$^{Y39TAG}$ knock-in construct, together with the NES PylRS/tRNA$_{CUA}^{Pyl}$ and eRF1$^{E55D}$. After incubation for 6 days with or without TCO*A-Lys, neurons were fixed and endogenous Tubb3 tagged with GFP$^{Y39\rightarrow UAA}$ was imaged with widefield microscopy. Scale bars: 50 µm (**a**,**b**).

**Supplementary Tables**

**Supplementary Table 1.** Primers used for cloning and mutagenesis

**Supplementary Table 2.** Background values used for the deconvolution of confocal and STED images

**Supplementary Table 1.** Primers used for cloning and mutagenesis

| Purpose | Primer name | Primer sequence 5'–3' |
|---|---|---|
| Cloning of NFL in mEGFP-N1 plasmid | NfL_HindIII_fw | GGT GGT AGC TTC ACC ATG AGT TCG TTC GGC TAC GAT |
| | NfL_ApaI_rv | ACC ACC GGG CCC CAT CTT TCT TCT TAG CCA CCT GCT CC |
| Mutagenesis of NFL, position K211 | NFL_K211TAG_fw | CTG GAG TAG CGC ATC GAC AGC CTG ATG GAC |
| | NFL_K211TAG_rv | GAT GCG CTA CTC CAG CTC GGC GCG |
| Mutagenesis of NFL, position K363 | NFL_K363TAG_fw | AGC ACG TAG AGC GAG ATG GCC AGG TAC CT |
| | NFL_K363TAG_rv | CTC GCT CTA CGT GCT TCT CAG CTC ATT CTC CAG T |
| Mutagenesis of NFL, position R438 | NFL_R438TAG_fw | TCT GCT TAG TCT TTC CCA GCC TAC TAT ACC AGC CA |
| | NFL_R438TAG_rv | GAA AGA CTA AGC AGA CAT CAA GTA GGA GCT GCT |
| Mutagenesis of NFL, position K468 | NFL_K468TAG_fw | GAG GCC TAG GAT GAG CCC CCC TCT GAA GGA |
| | NFL_K468TAG_rv | CTC ATC CTA GGC CTC CTC AGC TTT CGT AGC |
| Cloning of $NFL^{WT}$-FLAG and $NFL^{TAG}$-FLAG | BamHI_FLAG_NotI_fw | GA TCC G GAC TAC AAA GAC GAT GAC GAC AAG TGA GC |
| | BamHI_FLAG_NotI_rv | GGC CGC TCA CTT GTC GTC ATC GTC TTT GTA GTC CG |
| Cloning of U6-$tRNA_{CUA}^{Pyl}$ in pcDNA3.1/Zeo(+)_NES $PylRS^{AF}$ plasmid | U6-tRNAcassette_BglIIfw | GGTGGT AGATCT AAA AAA CGG AAA CCC CGG GAA TCT AAC C |
| | startU6cassette_MfeIrv | CCC TTT CAA TTG GAG GGC CTA TTT CCC ATG ATT CCT TCA TAT TTG |
| Cloning of pORANGE Tubb3-$GFP^{Y39TAG}$ KI | Tubb3-GFP-KI_HindIII_fw | GGA GGA AAG CTT GCT GCG AGC AAC TTC ACT TGG GGG ATC AGG CGT GAG CAA GGG CGA GGA GC |
| | Tubb3-GFP-KI_XhoI_rv | TCC TCC CTC GAG CCC AAG TGA AGT TGC TCG CAG CAC ATT ACT TGT ACA GCT CGT CCA TGC CGA G |
| Cloning of NFL target sequence into the pORANGE cloning template vector | NFL_gRNA_BbsI_fw | CACC GAG TGC TGG AGA GGA GCA GG |
| | NFL_gRNA_BbsI_rv | AAA C CCT GCT CCT CTC CAG CAC TC |
| Cloning of linker-3xFLAG and $linker^{A6TAG}$-3xFLAG donor sequences in pORANGE NfL KI plasmid | NFL-link-3xFLAG KI_HindIII_fw | GGA GGA AAG CTT CC ACC TGC TCC TCT CCA GCA CTC GGT AGC GCT GGA AGC GCT |
| | NFL-link(A6)-3xFLAG KI_HindIII_fw | GGA GGA AAG CTT CC ACC TGC TCC TCT CCA GCA CTC GGT AGC GCT GGA AGC TAG GAC TAC |
| | NFL-link-3xFLAG KI_BamHI_rv | TCC TCC GGA TCC GAG TGC TGG AGA GGA GCA GGT GGT TAC TTG TCA TCG TCA TCC TTG TAA TCG ATG TCA TG |

**Supplementary Table 2.** Background values used for the deconvolution of confocal and STED images

| Figure | Image | Background value |
|---|---|---|
| Fig. 4a | SiR-tz confocal | 5 |
| | SiR-tz STED | 0.5 |
| Fig. 4b | SiR-tz confocal | 4 |
| | SiR-tz STED | 5 |
| | ATTO488-tz confocal | 3 |
| | ATTO488-tz STED | 5 |
| Fig. 5b | SiR-tz STED | 1 |
| | ATTO488-tz STED | 4 |
| Fig. 5c | SiR-tz STED | 3 |
| | ATTO488-tz STED | 4 |
| Supplementary Fig. 7a | ATTO488-tz confocal | 2.5 |
| | ATTO488-tz STED | 1.5 |
| Supplementary Fig. 7b | CF650-me-tz confocal | 5 |
| | CF650-me-tz STED | 1.5 |
| Supplementary Fig. 7c | SiR-tz confocal | 2 |
| | SiR-tz STED | 2 |
| Supplementary Fig. 8 | SiR-tz confocal | 5 |
| | SiR-tz STED | 2 |
| | ATTO488-tz confocal | 3 |
| | ATTO488-tz STED | 4 |
